# TuBA: Tunable Biclustering Algorithm Reveals Clinically Relevant Tumor Transcriptional Profiles in Breast Cancer

**DOI:** 10.1101/245712

**Authors:** Amartya Singh, Gyan Bhanot, Hossein Khiabanian

## Abstract

**Background:** Traditional clustering approaches for gene expression data are not well adapted to address the complexity and heterogeneity of tumors, where small sets of genes may be aberrantly co-expressed in specific subsets of tumors. Biclustering algorithms that perform local clustering on subsets of genes and conditions help address this problem. We propose a graph-based Tunable Biclustering Algorithm (TuBA) based on a novel pairwise proximity measure, examining the relationship of samples at the extremes of genes’ expression profiles to identify similarly altered signatures.

**Results:** TuBA’s predictions are consistent in 3,940 Breast Invasive Carcinoma (BRCA) samples from three independent sources, employing different technologies for measuring gene expression (RNASeq and Microarray). Over 60% of biclusters identified independently in each dataset had significant agreement in their gene sets, as well as similar clinical implications. About 50% of biclusters were enriched in the ER-/HER2- (or basal-like) subtype, while more than 50% were associated with transcriptionally active copy number changes. Biclusters representing gene co-expression patterns in stromal tissue were also identified in tumor specimens.

**Conclusion:** TuBA offers a simple biclustering method that can identify biologically relevant gene co-expression signatures not captured by traditional unsupervised clustering approaches. It complements biclustering approaches that are designed to identify constant or coherent submatrices in gene expression datasets, and outperforms them in identifying a multitude of altered transcriptional profiles that are associated with observed genomic heterogeneity of diseased states in breast cancer, both within and across tumor subtypes, a promising step in understanding disease heterogeneity, and a necessary first step in individualized therapy.

## BACKGROUND

The first step in organizing and analyzing high-throughput gene expression datasets is to group together (cluster) genes, or samples based on some mathematical measure of similarity between the respective entities of interest. Since a priori knowledge about both the relevant genes, and the unique phenotypic characteristics of samples is usually limited, clustering is often performed in an unsupervised manner [1–5]. Quite frequently, measures of similarity (such as the Pearson correlation coefficient, Spearman correlation coefficient, Mutual Information, etc.) are employed to quantify the level of similarity between every pair of genes (or samples) across all samples (or genes). Such an approach is known as global clustering. In case of datasets with a heterogeneous assortment of samples, only a small subset of genes in a fraction of the total set of samples may be co-regulated in specific cellular processes. This is especially true for diseases like cancer that manifest a plethora of diseased phenotypes. In gene expression datasets comprising tumor samples, depending on the heterogeneity of the diseased states, there may be multiple distinct transcriptional alterations that are exhibited by multiple (not necessarily exclusive) subsets of the tumor samples. Moreover, it is well known that even in normal cells, the same genes can regulate and participate in multiple distinct pathways, depending on the context. Therefore, global clustering is not an optimal approach to identify co-expressed sets of genes or samples in gene expression datasets associated with heterogeneous diseases.

To address these concerns, a variety of local biclustering algorithms have been proposed that satisfy the following requirements: 1) a cluster of genes is defined with respect to only a subset of conditions (patient samples) and vice versa, and 2) the clusters are not exclusive and/or exhaustive – i.e. a gene/condition may belong to more than one cluster or to none at all [6–9]. Based on the type of biclusters and the mathematical formulation used to discover them, biclustering techniques are categorized by Oghabian *et al* [10] into four classes: (i) correlation maximization methods that identify subsets of genes and samples where the expression values of genes (or samples) is highly correlated across samples (or genes) [6]; (ii) variance minimization methods that identify biclusters where the expression values have low variance among the selected genes or conditions or both [11]; (iii) two-way clustering methods that iteratively perform one-way clustering on the genes and samples [12], and (iv) probabilistic and generative methods that employ stochastic approaches to discover genes (or samples) that are similarly expressed in subsets of samples (or genes) [13, 14]. Another classification scheme proposed by Pontes et al [15, 16] categorizes the generated biclusters based on their gene expression patterns into four classes: (i) biclusters with constant values, (ii) biclusters with constant values on rows (genes) or columns (conditions), (iii) biclusters with additive and/or multiplicative relationships between genes and conditions, and (iv) biclusters based on evidence that a subset of genes is up-regulated or down-regulated across a subset of conditions without taking into account actual expression values; data in such biclusters does not follow any mathematical model.

In this paper, we introduce a graph-based method, called the Tunable Biclustering Algorithm (TuBA), which discovers biclusters consistent with the latter category. TuBA is based on a novel measure of proximity that identifies aberrantly co-expressed gene sets within subsets of tumor samples that correspond to the expression extremals for the genes. A key feature of the proximity measure used in TuBA is that it does not rely explicitly on the actual gene expression values. We demonstrate the utility of TuBA by applying it to three large, independent cohorts of breast invasive carcinoma (BRCA) encompassing 3,940 patients. In addition to detecting known pathways and subtypes associated with breast cancer, TuBA was able to uncover several novel sets of co-expressed genes across subtypes that may be relevant as biomarkers for therapeutic identification and intervention.

## METHODS

### Proximity measure

TuBA’s proximity measure addresses the following question: in a given gene expression dataset, which genes exhibit higher (or lower) expression levels in the same subset of samples relative to the rest? In other words, if we only consider the top (or bottom) *x* percentile samples for every gene, which gene-pairs share a significant number of samples between their percentile sets? The number of samples shared between any pair of percentile sets follows the hypergeometric distribution; therefore, we can compute the significance (p-value) of overlaps between pairs of percentile sets based on the numbers of shared samples by using the one-sided Fisher’s exact test. Thus, TuBA’s proximity measure between two genes is defined by the significance of overlaps between their respective percentile sets (**Fig. 1**).

**Fig. 1.**
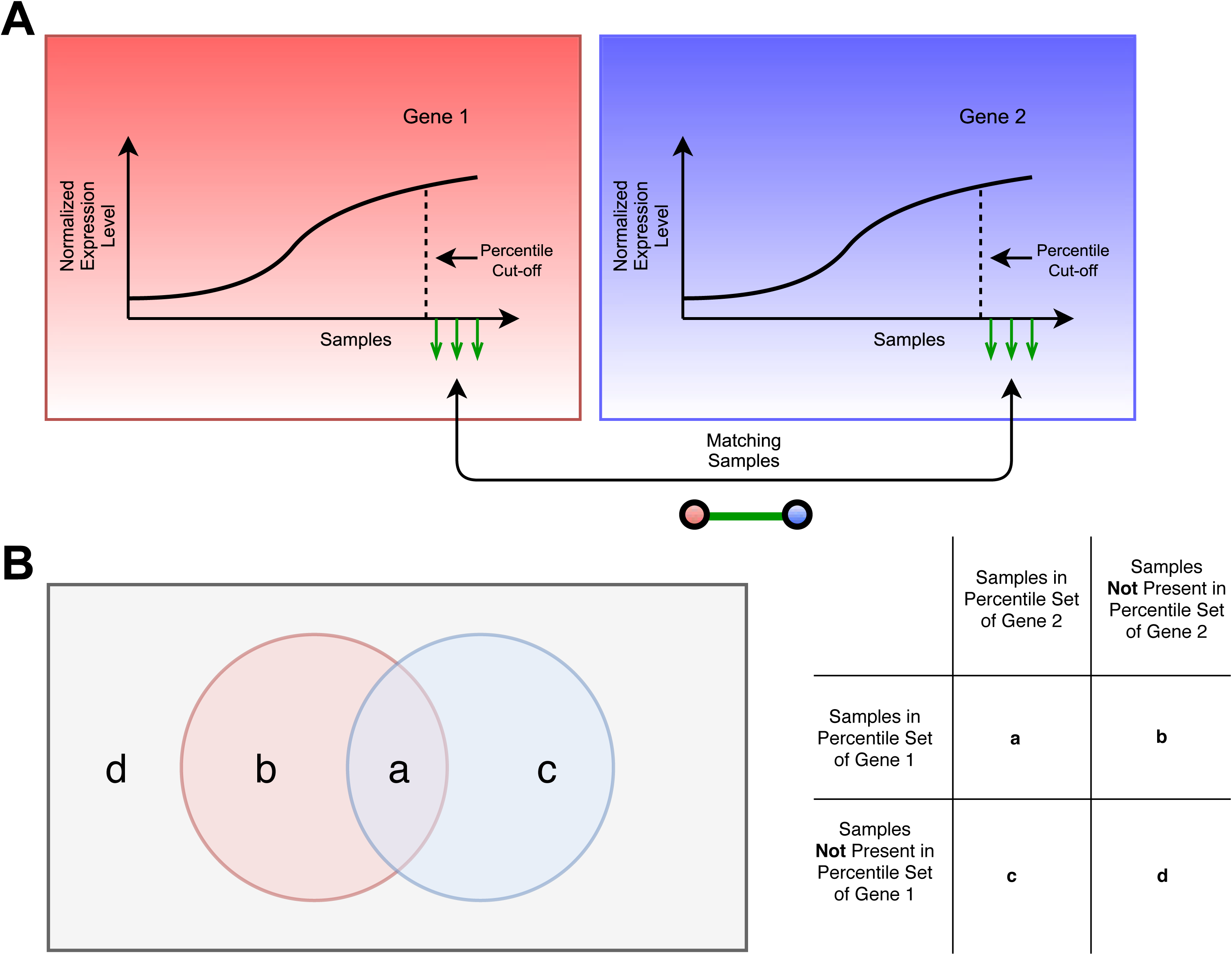
Schematic representation of TuBA’s proximity measure. (A) For each gene, samples are arranged in increasing order of expression levels and those corresponding to a fixed percentile set (top or bottom) are compared between each pair of genes as shown. The gene-pairs that share a significant number of samples are represented as nodes linked by edges, which represent the samples. (B) The Venn diagram illustrates the setup of the contingency table for the one-sided Fisher’s exact test. The grey rectangular box represents the set of all samples in the dataset, the red and blue circles represent the samples in the top (or bottom) percentile sets of gene 1 and gene 2, respectively.

In a real biological dataset, we expect the following two scenarios to arise: (i) subsets of genes associated with particular biological processes/pathways are co-expressed in all samples. In this case, it is reasonable to expect a significant agreement between the sets of samples that exhibit higher (or lower) expression levels of the involved genes. (ii) Alternatively, subsets of genes may be dysregulated via shared underlying mechanisms, such that their expression levels are higher (or lower) compared to the rest of the samples that are not influenced by that mechanism. The latter case is of particular interest for datasets associated with diseased states, especially cancers, since these gene co-expression signatures and their underlying mechanisms could help us identify potential biomarkers with prognostic and/or predictive value. This is the basic motivation behind standard different differential expression analyses as well [17]. However, unlike usual differential co-expression analyses, our proximity measure does not rely on any pre-specification of subtypes.

A salient feature of our proximity measure is that it does not model the distributions of the measured expression levels of genes across samples. Moreover, it does not rely on significant differences between the expression levels of genes in samples comprising the extremal sets versus the rest of the samples. Thus, biologically relevant gene co-expression signatures can be identified without restricting the analysis exclusively to genes that exhibit differential expression across subsets of samples. In case of tumor datasets, this increases the likelihood for identification of gene co-expression signatures associated with the microenvironment. Another salient feature of our proximity measure is that there is no penalty for relative changes in ranks of samples in the respective percentile sets. This is important because, even if the ranks of matching samples are significantly different in the two percentile sets, there is still valuable information to be gleaned by virtue of the fact that these subsets of samples exhibit higher (or lower) expression levels for a given gene-pair compared to all the other samples. This feature of our proximity measure makes it less sensitive to noise compared to other proximity measures, such as the Spearman’s Rank correlation.

### Graph-based Algorithm

For each gene, TuBA identifies samples in the upper-most (or lower-most) percentile sets. Pairwise comparison between these percentile sets (using the one-sided Fisher’s exact test) identifies gene-pairs that share a statistically significant number of samples. Each significant gene-pair is illustrated graphically as a pair of nodes connected by an edge that represents the samples shared between their percentile sets. The complete set of these pairwise graphs generates large graphs from which robust gene co-expression signatures are recovered using the following iterative process (**Fig. 2**):

1. The graph is pruned such that its elementary units are complete subgraphs (cliques) of size 3 (triangles).
2. The largest clique (i.e. the seed) in the pruned graph is identified using the Bron-Kerbosch algorithm [18]. In cases where the largest clique is not unique, the union of all equally large cliques with a non-zero intersection of their nodes is designated as the seed; the remaining largest cliques are identified as new seeds in subsequent iterations.
3. The graph is trimmed by removing all the edges that contain any of the nodes in the seed in step 2. This step significantly reduces the computation time required to identify all the robust cliques in the graph.
4. Steps 2 and 3 are repeated till the graph has no elementary units left.
5. The seeds identified in steps 1-4 are exclusive in their gene sets, i.e., no two seeds share a common gene. To create the bicluster, the seeds are reintroduced sequentially into the original pruned graph from step 2, and nodes that share edges with at least two nodes in each seed are identified and added to the seed. The resulting graphs are the final biclusters obtained by TuBA.

**Fig. 2.**
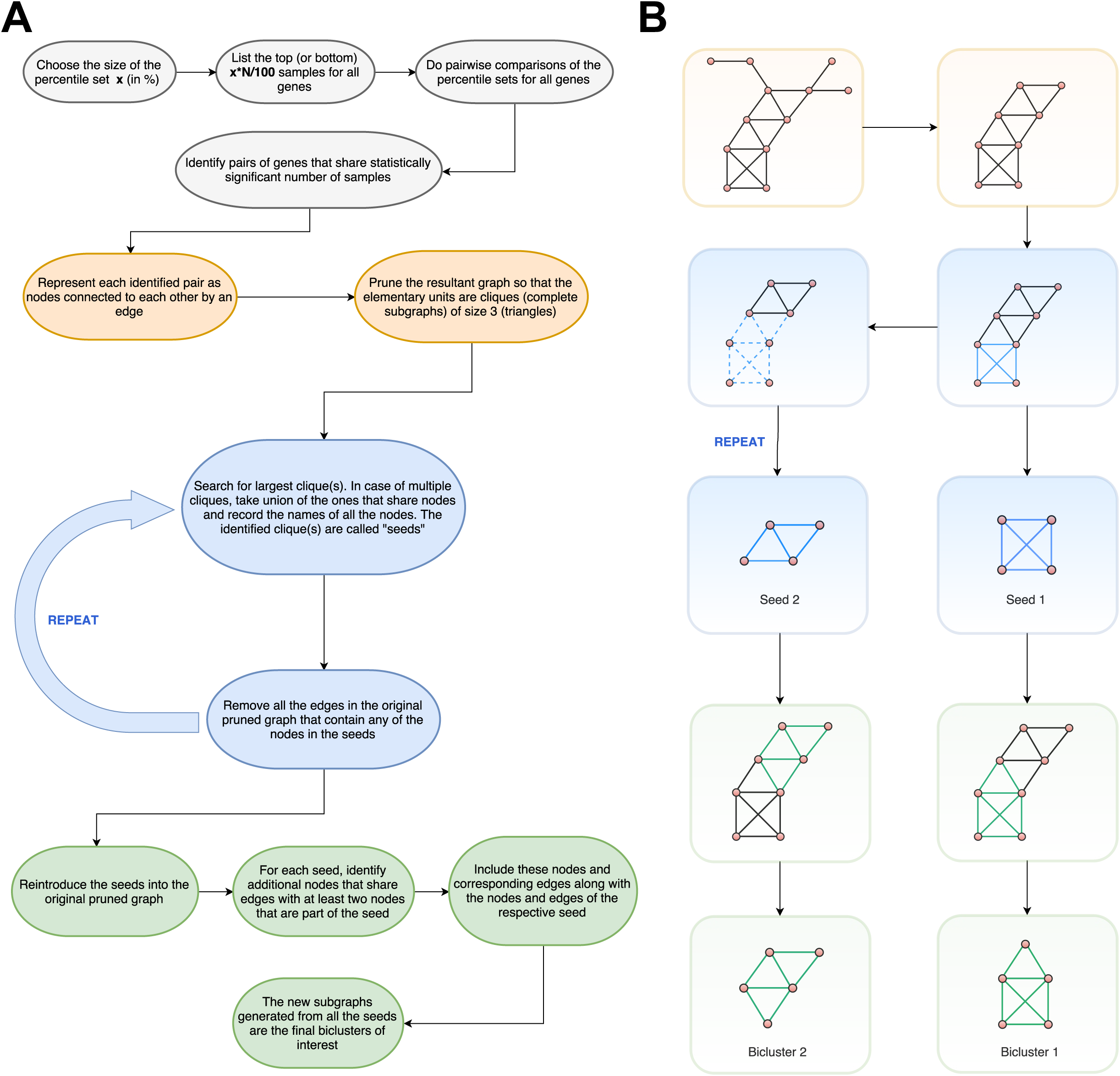
TuBA’s schematics. (A) Flowchart of the pipeline for TuBA. (B) Schematic representation of the graph-based approach to discover biclusters.

Note, the requirement of largest cliques as seeds of our biclusters in step 2 is a key step in our algorithm that enables the identification of shared altered mechanisms in subsets of samples that exhibit high (or low) expression levels of these genes, while permitting the study of sets of co-expressed genes that are associated with functionally related pathways. Implicit in this requirement is the crucial assumption that the sets of genes comprising the largest cliques are co-expressed in a subset of samples that comprise the edges. This assumption is not the same as requiring all gene-pairs comprising the seed to share identical sets of samples, or assuming that all the samples comprising the final biclusters co-express all the genes present in the bicluster. Instead, our expectation is that the samples present in the final biclusters are *enriched* in the top (or bottom) samples for each gene comprising the biclusters. We have provided supporting evidence for this expectation in the **Results** section.

Gene enrichment analysis of the gene sets in the biclusters can be used to identify their functional relevance, and sample enrichment analysis can elucidate potential clinical subtypes, underlying mechanisms of disease, and possible therapeutic approaches. Furthermore, within each bicluster, genes can be assigned degrees, which are the total number of edges that connect them to other genes in the graph. Genes with higher degrees exhibit co-expression with other genes in a greater proportion of samples in the bicluster. These could be candidate driver genes.

### Tuning TuBA

TuBA has two adjustable parameters:

1. The Percentile cutoff: This parameter controls the number of top or bottom samples (based on expression levels) considered for comparison between genes.
2. The Overlap significance cutoff: The p-value threshold used to assess significance of the overlap of samples between percentile sets for each gene-pair. This parameter controls the minimum number of samples that must be shared between percentile sets for an association to be considered significant, and to be represented in the graph.

It is best to interpret the parameters as “knobs” that can be tuned to probe different levels of heterogeneity in the population. For a given dataset, the choice of these two parameters determines the number, as well as the composition of the final biclusters. The choice of the first parameter determines the level of heterogeneity and/or the extent of prevalence of genomic alterations in tumors that may be of interest to the investigator. To illustrate how the choice of percentile cutoff could affect the identification of co-expressed gene pairs, we consider a hypothetical dataset consisting of 200 samples. Assume that there is a gene-pair in this dataset that is up regulated in 5% of the samples such that the top 5% percentile sets (i.e. top 10 samples) are identical for the two genes. **Fig. 3A** shows the significance values for overlaps (p-values calculated using one-sided Fisher’s exact test) as a function of the fraction of samples that overlap. At 5% percentile cutoff, the significance value for an overlap fraction of 1 (complete match/overlap between percentile sets) is *p* = 4.45e-17 (dark blue curve). If instead we had chosen an upper 10% percentile cutoff, we would have an overlap fraction of 0.5 (overlap of 10 out of 20 samples) corresponding to a significance value for overlap between 1e-10 and 1e-5. Thus, an increase in the size of the percentile set results in loss of significance for aberrant co-expression signatures found in smaller subsets of samples. This does not imply that it is generally better to choose smaller percentile sets. In fact, a reduction in the size of the percentile set increases the likelihood that a number of samples match purely by chance. (Note the p-values corresponding to the overlap fraction of 1 for the three cases in **Fig. 3A**, further demonstrated by permutation tests in Results.) Thus, as we vary the size of the percentile set, there exists a trade-off between the sensitivity (identification of altered transcriptional profiles in small subsets of population) on one hand and the overlap significance on the other.

**Fig. 3.**
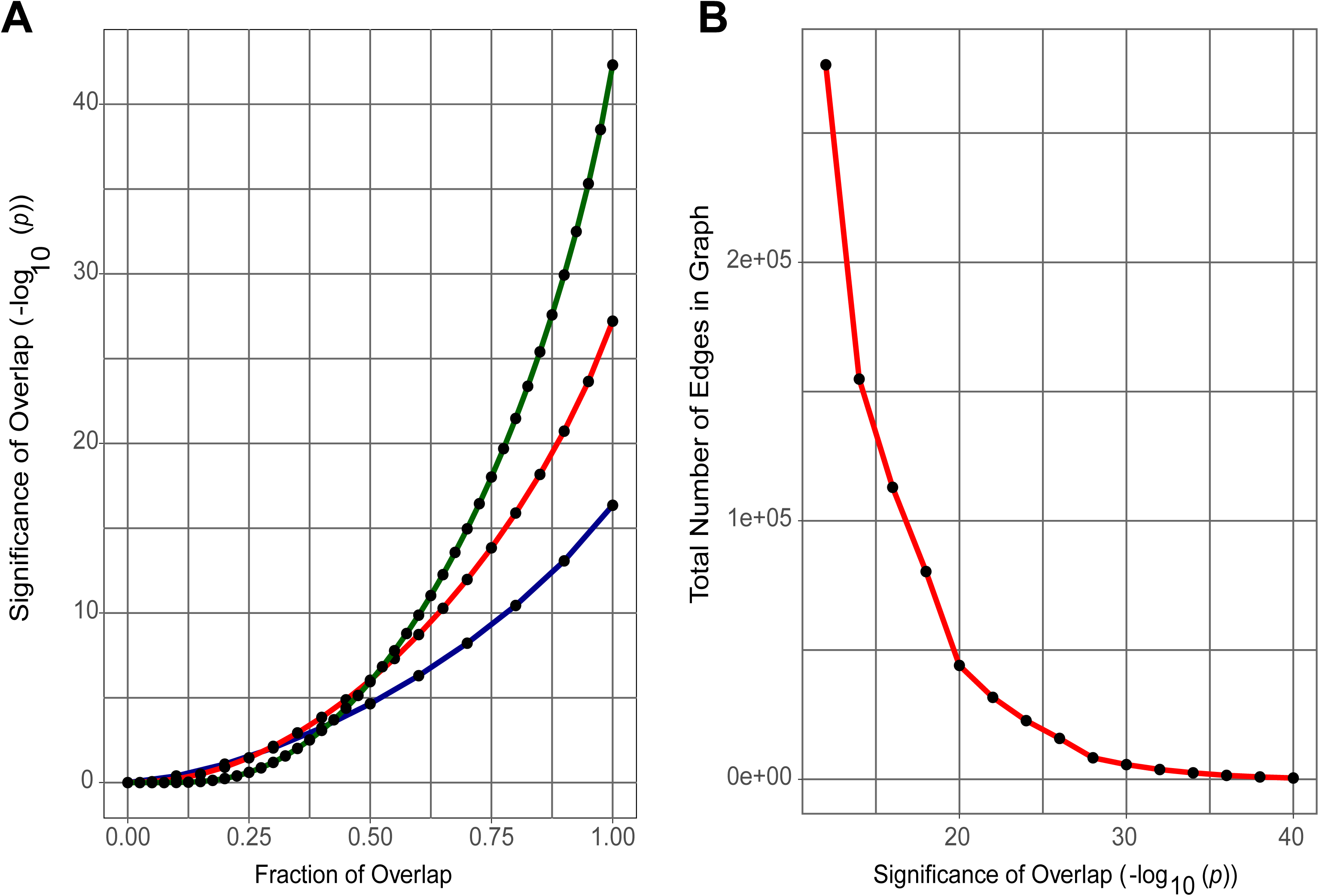
Tuning TuBA’s parameters. (A) Significance of overlap corresponding to fraction of overlap between 0 (no matches/overlap) & 1 (all samples match/overlap) for percentile set size of: (i) top 20% (green), (ii) top 10% (red), and (iii) top 5% (dark blue) respectively for a hypothetical dataset consisting of 200 samples. (B) Divergence of the total number of edges in the graph for the TCGA RFS dataset as we lower the cutoff for the significance of overlap.

The choice of the second parameter – the extent of patient/sample overlap between percentile sets – determines the gene-pairs that will be represented in the graph that is explored iteratively to identify sets of co-expressed genes. As we lower the significance of overlap (increasing p-values), new genes and samples get added to the graph resulting in an increase in the number of edges (**Fig. 3B**). Further lowering of the overlap significance results in the addition of many more edges to the graph, however this addition is not accompanied by a proportional increase in the samples or genes added to the graph. As more edges get added to the graph, the computational effort required for finding maximal cliques increases. Since the maximal clique problem is NP-hard [19], it can take exponential time to find all maximal cliques. Thus, the cutoff for the significance of overlap is informed by the trade-off between the gain of new information in the biclusters in terms of new samples and genes, and the number of edges added to the graph that leads to a disproportionate increase in the computational effort. We propose the following heuristic for choosing the cutoff value: the cutoff for the significance level of overlap should be such that a decrease in the significance level by an order of magnitude leads to an 40-60% increase in the number of edges that get added to the graph (Note the number of edges that get added to the graph at overlap significance p-values larger than 1e-20 in **Fig. 3B**).

Because the principal goal of TuBA is to identify subsets of genes that are co-expressed at high (or low) levels within subsets of samples, the exact number of biclusters is not biologically relevant despite possible small variations in their total number as the algorithm is tuned. We investigated the consistency of TuBA’s biclusters across different choices of the parameters. We used the hypergeometric test to identify biclusters that share significant fractions of their genes, and observed that despite a five-fold difference in the significance level of overlap, there is greater than 80% agreement between the sets of biclusters obtained for different choices of the overlap significance cutoff. The results of the analysis are presented and discussed under **Robustness of TuBA’s Biclusters** in **Supplementary Methods** and **Supplementary Table 4**.

### Datasets

We applied TuBA to three independent BRCA datasets that employed distinct methods for measuring transcript levels: (i) TCGA RNASeq gene expression dataset using the Illumina HiSeq 2000 RNA sequencing platform, (ii) METABRIC gene expression dataset using the Illumina HT-12 v3 microarray platform, and (iii) six cohorts with gene expression data from GEO using the Affymetrix HGU133A microarray platform. To compare results among datasets, we applied TuBA to only their common gene sets. For clinical association analysis, we prepared two separate datasets for patients with known recurrence free survival (RFS) status (908 patients) and patients with known PAM50 subtype annotation (522 patients) respectively. Henceforth, we will refer to TCGA RFS, METABRIC RFS and GEO RFS datasets simply as TCGA, METABRIC and GEO, respectively, and PAM50 datasets are indicated specifically.

1. **TCGA – BRCA:** The log_2_(x+1) transformed RSEM normalized counts of Level 3 data (2016-08-16 version), the clinical data (including relapse status and PAM50 subtype annotation from the 2012 Nature study [20]) (2016-04-27 version), and gene-level copy number variation (CNV) data, as estimated by GISTIC2 [21] (2016-08-16 version) were downloaded from the UCSC Xena Portal (http://xena.ucsc.edu). Genes with zero expression in all samples as well as the samples with NAs for any gene were removed from the analysis.
2. **METABRIC:** Normalized gene expression data, the clinical file, and the copy number file for the METABRIC study were downloaded from the cBioPortal (http://www.cbioportal.org) on 2017-05-14 [22, 23]. Gene expression dataset of 1,970 samples that had both relapse status and PAM50 subtype annotation were used in this study.
3. **GEO:** MAS5 normalized gene expression data and the clinical data with relapse status were downloaded from [24] on 2017-05-10. The dataset comprises samples from six independent cohorts. After processing, our gene expression dataset consisted of 1,062 patients with relapse status.
4. **breastCancerNKI (Bicmix):** Gene expression data from breast cancer study published by van’t Veer et al. in 2002 [25], and van de Vijver et al. in 2002 [26] was downloaded in the form of an eSet using the breastCancerNKI package [27] in R. The dataset was further processed by removing probes with > 10% missing values, and imputing the missing values for the included probes [28].
5. **ESTIMATE:** Scores for the level of stromal cells present and the infiltration level of immune cells in tumor tissues for 906 out of 908 samples for the TCGA – BRCA RNA-SeqV2 dataset using the ESTIMATE algorithm were downloaded from http://bioinformatics.mdanderson.org/estimate on 2017-10-12.
6. **GTEx:** RNASeq raw counts data from the Genotype-Tissue Expression (GTEx) portal (www.gtexportal.org/home/) was downloaded on 2017-06-15. The dataset comprised of all tissue samples currently available in the GTEx database. The 214 breast tissue samples were identified and normalized using the DESeq package in R [29].

### Statistical Analysis

All computations were performed with R 3.3.0 [30]. The igraph package [31] was used to perform network/graph computations with some data summary functions performed using the plyr package [32]. The figures with graphs showing the genes for some of the biclusters were generated using Cytoscape v3.4.0 [33]. Permutation test was performed on the METABRIC dataset (1970 samples) with upper percentile set size cutoff: 5%. For each gene, we permuted the labels of the samples prior to ascertainment of the samples that corresponded to the top 5% respectively. The significance values for overlaps between every pair of genes were computed using the Fisher’s exact test. We performed 100 iterations of these permutation tests in total. The data.table package was used to handle data files, and the ggplot2 package was used to make plots [34]. GeneSCF [35] was used to perform gene set clustering based on functional annotation and to associate biclusters with specific biological processes. A binary matrix with biclusters along the rows and samples along the columns was generated to perform hierarchical clustering. Samples belonging to respective biclusters were assigned a value of 1. The Hamming distance was used to measure dissimilarity between the biclusters as well as the samples. All tests for enrichments were done using the Fisher’s exact test; where necessary the p-values were corrected for multiple hypotheses testing using the Benjamini–Hochberg false discovery rate method [36]. The details of the contingency tables for the tests are provided in **Supplementary Methods**. All enrichments are reported at FDR < 0.05, unless specified otherwise.

## RESULTS

### Benchmarking TuBA’s Proximity Measure

TuBA’s pairwise proximity measure for genes is based on overlaps between their extremal subsets of samples. In comparison, global pairwise linear correlation coefficients such as the Pearson’s correlation coefficient and Spearman’s rank correlation coefficient are both susceptible to the influence of outliers. Even in the absence of outliers, genes that are co-expressed across all samples are expected to have significant overlaps between the subsets of samples that correspond to their top (or bottom) percentile sets. Given these observations, we tested the hypothesis that gene sets identified by global proximity measures have significant overlap with those identified by TuBA within its biclusters.

We computed the Pearson’s correlation coefficients between all possible pairs of genes in the TCGA and METABRIC datasets, respectively. We shortlisted all gene-pairs with the correlation coefficient greater than or equal to 0.6. We then employed our graph-based algorithm to identify gene co-expression modules within the graphs. For TCGA and METABRIC, we obtained 569 and 298 gene co-expression modules, respectively (**Supplementary Table 1**). We investigated the association between the gene sets in biclusters discovered by TuBA’s proximity measure versus gene co-expression modules identified by global correlation metrics by performing a hypergeometric test for gene overlaps. The null hypothesis was the absence of significant overlap between their respective sets of genes. More than 89% (316 out of 353 biclusters) of the biclusters discovered by TuBA in the TCGA dataset comprised gene sets that were enriched in at least one gene co-expression module (FDR < 0.001), while 86% (293 out of 340 biclusters) of the biclusters discovered by TuBA in the METABRIC dataset were enriched in at least one module.

We performed a similar analysis using Spearman’s rank correlation, with a cutoff of 0.6 for the correlation coefficient. We obtained 524 and 232 gene co-expression modules for TCGA and METABRIC, respectively. More than 80% (285 out of 353 biclusters) of TuBA’s biclusters in the TCGA dataset comprised gene sets that were enriched in at least one gene co-expression module (FDR < 0.001), while 73% (249 out of 340 biclusters) of the biclusters discovered by TuBA in the METABRIC dataset were enriched in at least one module. Overall, we see a significant enrichment of gene sets in TuBA’s biclusters with the co-expression modules obtained by using the two global proximity measures. This is in concordance with our hypothesis stated earlier.

As noted earlier, due to samples that exhibit aberrant/outlier expression of some genes, the linear correlation coefficients can often get skewed to reflect greater pairwise correlations between such genes that our graph-based algorithm can identify. However, due to the global nature of these proximity measures, the resulting graphs lack any information on the samples that might be associated with aberrant expression of these genes. In other words, unlike the case for TuBA, the edges in these graphs do not represent any subset of samples; they simply reflect an association between the genes by virtue of their pairwise correlation coefficient being greater than the chosen cutoff. The novel design of our proximity measure enables precise identification of co-expressed gene sets, while discerning the subsets of samples that exhibit higher (or lower) expression levels of these genes relative to the rest of the samples.

### Choice of parameters for TuBA

For a given choice of the size of percentile set, TuBA generates plots that illustrate the number of added genes, added edges, and added samples as the overlap significance cutoff is varied. These are used to inform the choice of the overlap significance cutoff based on our proposed heuristic. Since the experimental platform and the total number of samples were different among the analyzed datasets, the choice of the overlap significance cutoff varied (**Fig. 4**). For the respective choices of the knobs, we obtained 353, 340, and 369 biclusters for the TCGA, METABRIC, and GEO datasets, respectively (**Supplementary Table 2**). Permutation tests showed that no gene-pairs had p-values less than the cutoffs (**Fig. S1**). Moreover, after adjusting for multiple hypotheses testing, none of the gene-pair p-values were statistically significant.

**Fig. 4.**
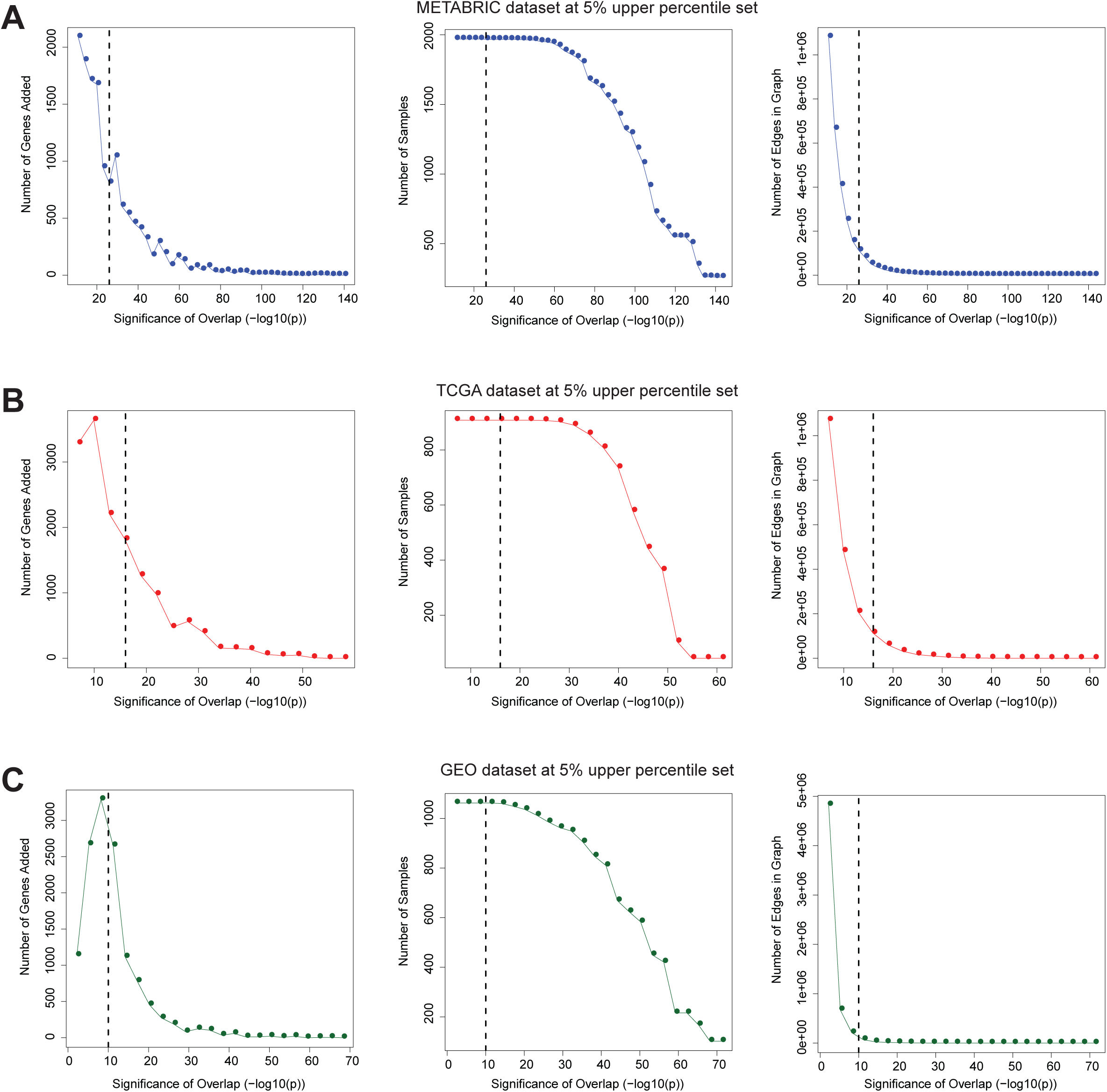
The effect of TuBA’s parameters on the number of genes and samples in the graph. Plots for the number of genes added to the graph for every incremental decrease in the significance level for overlap (−log_10_(*p*)), the number of samples in graph at different significance levels of overlap and the total number of edges in graph at different significance levels of overlap corresponding to a percentile set size of 5% for (A) METABRIC, (B) TCGA, and (C) GEO datasets, respectively.

### Enrichment of bicluster samples in top (bottom) sample sets of bicluster genes

As pointed out earlier, the identification of largest cliques as seeds of our biclusters was based on the expectation that samples present in the final biclusters were enriched specifically in the up (down)-regulated samples for each gene comprising the cliques. We tested the hypothesis that the subsets of samples comprising the biclusters were enriched by the samples that comprise the top (bottom) sets for each gene in the bicluster. For example, suppose a dataset consists of 1000 samples, wherein on application of TuBA for high expression, one bicluster is identified to comprise 100 genes and 200 samples. For each of those 100 genes, we identify their top 200 samples and test whether these 200 samples are enriched in the 200 samples comprising the bicluster. The null hypothesis is that these two sets of samples are independent, and therefore we should not expect to see statistically significant associations between them. We applied this test for each of TuBA’s biclusters in the TCGA, METABRIC, and GEO datasets. For high expression, we observed that all genes in all 353 biclusters from TCGA showed significant enrichment (hypergeometric test FDR < 0.001). In case of METABRIC we observed two biclusters (bicluster 2 and bicluster 269) out of 340 biclusters with only 1% of their constituent genes not exhibiting enrichment, while in case of GEO we observed only one bicluster (bicluster 22) out of 369 with 1% of its constituent genes not exhibiting enrichment. For the low expression analysis of TCGA, we observed two biclusters (bicluster 14 and bicluster 165) out of 203 biclusters that comprised a few genes that did not exhibit enrichment. A crucial observation across all the datasets was that even in the few biclusters that included a few genes with enrichment FDR greater than 0.001, none of these genes were constituents of the seeds of those biclusters. We therefore found it justified to rely on the subsets of genes that comprise the seeds for future gene ontological enrichment tests. This enables us to identify the core functional signatures of biclusters.

We performed a similar analysis for the bicluster samples. Following the previous example of a high-expression bicluster with 100 genes and 200 samples, we aimed to identify the genes in the bicluster that had a given sample in their top 200 samples (based on the expression levels of the genes). Therefore, for each sample, we evaluated whether their corresponding subset of genes had significant overlaps with the complete set of genes comprising the bicluster. So, we tested the null hypothesis that overlaps between them were not statistically significant. For high expression, we observed that 95% of biclusters (336 out of 353) from the TCGA, 97% of biclusters (329 out of 340) in the METABRIC, and 89% of biclusters (328 out of 369) in the GEO databases had > 95% of samples enriched (hypergeometric test FDR < 0.001). For the low expression analysis of TCGA, we observed that 98% of biclusters (199 out of 203) had 95% of their samples enriched in the bottom 200 samples for the corresponding genes in the biclusters. Based on these analyses, the FDR values for each gene (sample) in any given bicluster can be viewed as their scores – the closer the value of the FDR is to zero for a gene (sample), the stronger is the association of the gene (sample) to the bicluster.

### Consistency of TuBA within a dataset

To investigate whether TuBA could consistently discover biclusters within the TCGA RFS cohort, the 908 samples were divided randomly into two groups of 454 samples each. This was done five times to generate five pairs of datasets. TuBA was applied to each dataset pair using a percentile set size of 5% and an overlap significance cut-off of FDR ≤ 1e-08. Pairwise comparisons (between sets of genes) of biclusters from the five trials showed that on average 73% biclusters from one dataset in each pair were enriched (FDR < 0.001) in at least one bicluster from the other (**Supplementary Table 3**). We found a significant difference (Mann-Whitney U-test *p* < 1e-05) in the number of genes contained in biclusters that matched among trials, compared to the number of genes in biclusters that did not; while the median size of biclusters that matched was 20 (range: 3–840), the median size of biclusters that did not match was 3 (range: 3–18) (note that 3 is the smallest-sized bicluster generated by TuBA). Overall, TuBA was able to consistently identify matching sets of co-expressed genes from randomly sampled subsets of data within a dataset.

### Consistency of TuBA’s biclusters among independent datasets

Using common sets of genes, we compared the biclusters obtained from: (i) TCGA and METABRIC, (ii) TCGA and GEO, and (iii) METABRIC and GEO. Pairwise comparisons of biclusters obtained from the two datasets were used to identify the biclusters that shared a significant proportion of their genes (FDR < 0.001). In the TCGA vs. METABRIC comparison, 64% of biclusters obtained in one dataset were enriched in at least one bicluster in the other. In the TCGA vs. GEO comparison, 69% of biclusters obtained in one dataset were enriched in at least one bicluster in the other. Finally, in the METABRIC vs. GEO comparison, 76% of the biclusters obtained in one dataset were enriched in at least one bicluster in the other. Once again, we found that the biclusters that did not match were significantly smaller (median number of genes: 3-5) than the biclusters that matched (median number of genes: 20-25) between the datasets (Mann-Whitney U-test *p* < 0.001).

### TuBA identifies subtype-specific biclusters

We classified BRCA samples based on the expression levels of the ESR1 (ER) and ERBB2 (HER2) genes into four subtypes: (i) ER-/HER2-, (ii) ER+/HER2-, (iii) ER-/HER2+, and (iv) ER+/HER2+ (where + corresponds to over expressed and – corresponds to under expressed). A substantial proportion of biclusters were enriched in the ER-/HER2-subtype – 53% for METABRIC (**Fig. 5A and 5B**), 54% for TCGA (**Fig. 5C and Fig. 5D**), and 40% for GEO (**Fig. S6**) (**Supplementary Table 5**).

**Fig. 5.**
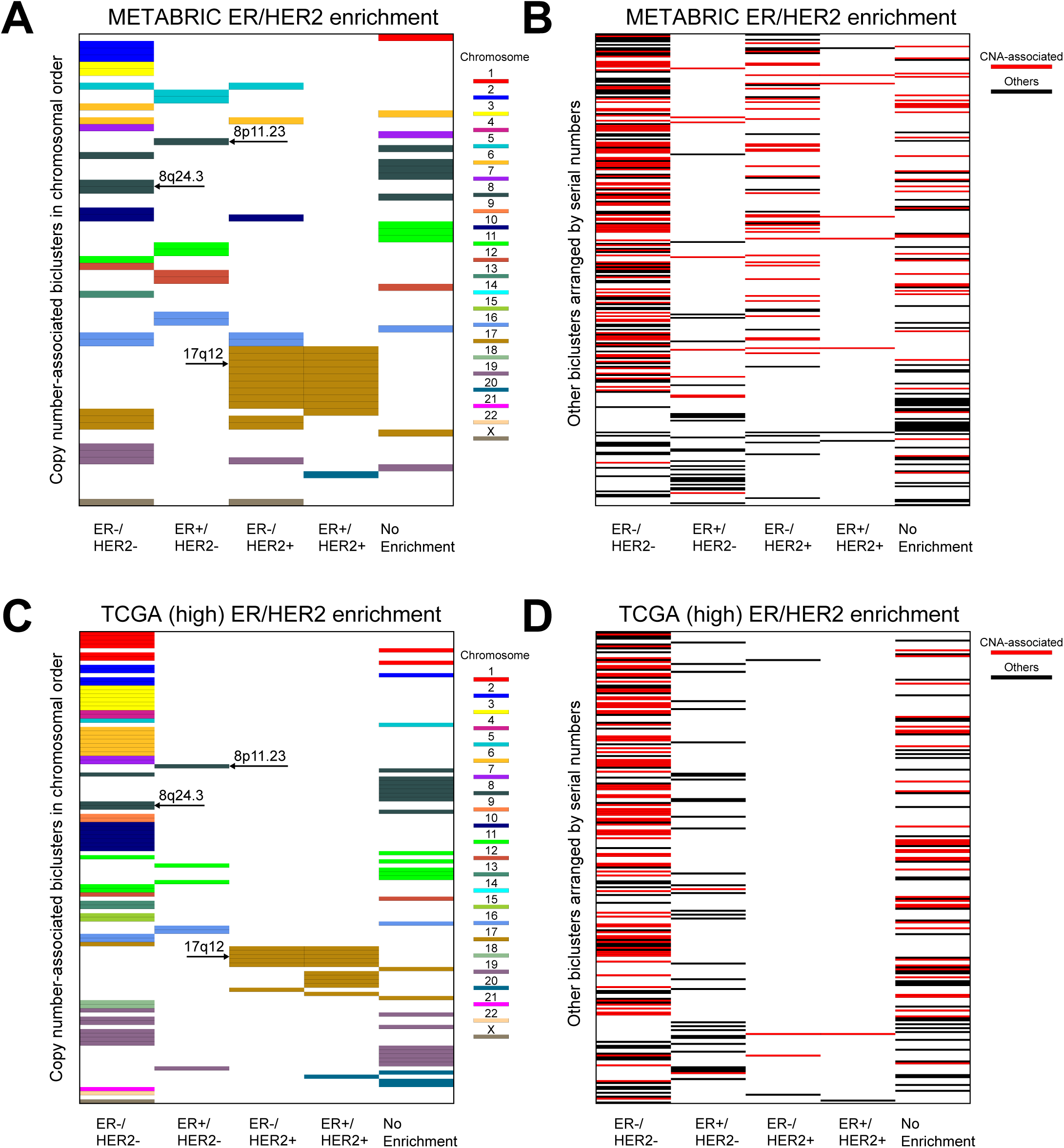
Subtype enrichment of CNV-associated and non-CNV biclusters. Enrichment of biclusters consisting of proximally located genes with copy number gains in the four subtypes based on ER/HER2 status for (A) METABRIC and (C) TCGA, respectively. The biclusters are represented by horizontal bars in each panel, color-coded according to the chromosome number of their constituent genes. Panels (B) and (D) show the remaining biclusters arranged according to their serial numbers in Supplementary Table 4 for METABRIC and TCGA, respectively. The ones that are associated with copy number (CN) gains of genes located at distant chromosomal sites are shown in red, while the rest are shown in black. Note, the thickness of the bar in each figure depends on the total number of biclusters displayed in that figure and so does not represent its chromosomal extent.

According to the PAM50 classification, there are five subtypes of BRCA: (i) Basal-like, (ii) Her2-enriched, (iii) Luminal A, (iv) Luminal B, and (v) Normal-like [37]. We observed a significant fraction of biclusters enriched in the basal-like subtype – 52% for METABRIC (**Fig. S4A and S4B**) and 55% for TCGA PAM50 (**Fig. S4C and S4D**). Although tumors of the basal-like or triple negative subtypes accounted for only about 15% of all BRCAs in the population, most of the altered expression profiles captured by our biclusters were in tumors of this subtype.

### TuBA identifies down-regulated subtype-specific biclusters in RNA-seq data

RNA sequencing offers a significant advantage over microarray assays. Theoretically, only the depth of sequencing limits the dynamic range of RNA-seq data [38, 39]. Given that TCGA’s RNA-seq data has adequate sequencing depth, we expected a reliable quantification of even lowly expressed transcripts. We therefore applied TuBA to the TCGA datasets to explore transcriptional profiles associated with low expression. We found that 46% biclusters from TCGA were enriched in the ER-/HER2-subtype (**Fig. S5A and S5B**), while 48% biclusters from the TCGA PAM50 dataset were enriched in the basal-like subtype (**Fig. S4E and S4F**). Thus, biclusters associated with low expression were predominantly enriched in the ER-/HER2- or basal-like subtypes. This further underscores the tremendous heterogeneity of altered transcriptional profiles within tumors of this subtype.

### TuBA highlights biclusters with proximally located genes

We observed that several biclusters discovered by TuBA across the three datasets comprise genes that are proximally located on the chromosomes, suggesting copy number amplification (CNA) as an underlying mechanism. Copy number data was used to calculate the significance of the proportion of samples present in each bicluster that exhibited copy number gains. For each gene in a given bicluster, we computed a p-value for the significance of the proportion of samples with CNA present in the bicluster. These p-values were then combined using Fisher’s method to yield a single p-value for the bicluster. This showed that 56% and 64% of biclusters from the METABRIC and TCGA datasets respectively were enriched for CNA (FDR < 0.001). Closer scrutiny revealed that only 60 (18%) biclusters from METABRIC were associated exclusively with CNA of proximally located genes (**Fig. 4A**), the remaining biclusters associated with CNA were enriched in genes from distant chromosomal locations (**Fig. 4B**). Similarly, 112 (32%) biclusters from TCGA were associated with CNA of proximally located genes (**Fig. 4C**). Many of these biclusters were associated with loci previously identified to exhibit copy number gains in BRCA [40, 41]. In order to explore the association between the biclusters obtained from the low expression analysis and loss of copy number, we repeated the copy number analysis described above. We observed that 52% biclusters from the TCGA dataset were enriched in copy number losses. However, only 21 biclusters contained genes located on the same chromosome (**Fig. S5A**), the remaining biclusters associated with copy number loss were enriched in genes from distant chromosomal locations. Similar analyses for PAM50 subtype enrichment for METABRIC and TCGA are summarized in **Fig. S4**.

To compare CNA associated biclusters between TCGA and METABRIC, we prepared two datasets that contained genes that were common in the two cohorts (17,209 genes). Pairwise comparison of the set of genes in the CNA enriched biclusters between the two datasets revealed that ~61% of biclusters from TCGA matched at least one CNA associated bicluster from METABRIC. On the other hand, 91% of biclusters from METABRIC were enriched in at least one CNA associated bicluster from TCGA. This suggests that most of the CNA enriched biclusters identified in the METABRIC microarray dataset were independently identified in the RNASeq dataset of TCGA.

We also observed some biclusters with proximally located genes that were not associated with gain in copy number. For TCGA, 14 biclusters out of 353 consisted of genes located proximally, while 18 biclusters out of 340 for METABRIC consisted of genes located near each other. Details of the genes and subtype-specific enrichments for some of these biclusters are summarized in **Supplementary Table 6**. Examples of biclusters from this category include the biclusters consisting of genes from the Cancer-Testis antigens family – *MAGEA2*, *MAGEA3*, *MAGEA6*, *MAGEA10*, *CSAG1*, *CSAG2*, *CSAG3* (Xq28)/*CT45A3*, *CT45A5*, *CT45A6* (Xq26.3). These genes are known to be aberrantly expressed in triple negative breast tumors [42], as well as in a few other tumor types [43].

### TuBA identifies biclusters associated with non-tumor expression signatures

We also discovered biclusters that appeared to be associated with non-tumor cells. For instance, biclusters associated with immune response were among the largest identified independently in all three datasets. The top five Gene Ontology – Biological Processes (GO-BP) terms for the bicluster associated with immune response were: T cell co-stimulation, T cell receptor signaling pathway, T cell activation, regulation of immune response, and positive regulation of T cell proliferation (**Supplementary Table 7**). This indicates immune cell infiltration in a significant number of tumor samples. To corroborate this, we stratified TCGA samples based on their ESTIMATE [44] scores for the infiltration level of immune cells in tumor tissues into three groups – (i) top 25 percentile, (ii) intermediate 50 percentile, and (iii) bottom 25 percentile – and verified that samples in these biclusters associated with immune response were enriched in samples with the highest levels of immune infiltration (FDR < 0.001)

For all three datasets, we also observed a bicluster associated with the stromal adipose tissue. The top 5 GO-BP terms for this bicluster were: response to glucose, triglyceride biosynthetic process, triglyceride catabolic process, retinoid metabolic process, and retinol metabolic process. An analysis based on the ESTIMATE scores for the level of stromal cells present in tumor tissue of TCGA samples confirmed that this bicluster was enriched within the top 25 percentile samples for stromal cell level. Subtype enrichment revealed that the bicluster was enriched in ER-/HER2-, basal-like (PAM50), and normal-like (PAM50) subtypes.

TuBA’s proximity measure was applied to gene expression data from 214 normal breast tissue samples from the Genotype-Tissue Expression (GTEx) public dataset. We observed that only 6.75% of biclusters obtained for the TCGA versus GTEx comparison were enriched in gene-pair associations identified in the GTEx dataset. The bicluster associated with the adipose tissue signature was one of the biclusters found enriched in GTEx. Another group of biclusters enriched in the three cancer datasets as well as in GTEx, were those associated with translation and ribosomal assembly. The top 5 GO-BP terms for these biclusters were: translation, rRNA processing, ribosomal small subunit biogenesis, ribosomal large subunit assembly, and ribosomal large subunit biogenesis. These biclusters were enriched in the ER-/HER2-subtype (FDR < 0.001).

### TuBA identifies clinically pertinent biclusters

We performed a Kaplan-Meier (KM) analysis of recurrence free survival (RFS), comparing the patients present in each bicluster to the rest for METABRIC and GEO. (The number of patients with incidence of recurrence in TCGA was insufficient for this kind of survival analysis to be statistically robust.) As expected for METABRIC, patients in the bicluster (bicluster 25) associated with the HER2 amplicon (17q12) had significantly shorter RFS time compared to the rest (**Fig. S7**). This is because patients in the METABRIC study were enrolled before the general availability of trastuzumab [45].

We also observed biclusters associated with CNA at the 8q24.3 locus in all three datasets (TCGA – biclusters 39 and 113, METABRIC – biclusters 26, 56, and 167, GEO – biclusters 16, 24, 37, 55, 74, 118, and 302). These patients also had significantly shorter RFS times compared to those patients whose tumors did not have amplification of this locus (**Fig. 6A**, **6B** and **6C**). A similar result was obtained when we restricted the samples to ER+/HER2-tumors, validating an earlier observation that copy number gain of the 8q24.3 locus may confer resistance to ER targeted therapy [46]. We note, however, that biclusters with amplification of the 8q24.3 locus were enriched in the ER-/HER2-subtype (*p* < 0.001). Hence, amplification of this locus may be even more relevant in determining treatment for patients with ER-/HER2-breast cancers assigned into an intermediate (ambiguous) risk class by Oncotype DX [46]. Genes at 8q24.3 that may be considered promising candidates based on their degrees in the biclusters include *PUF60*, *EXOSC4*, *COMMD5*, and *HSF1*. Specifically, *PUF60* is an RNA-binding protein known to contribute to tumor progression by enabling increased *MYC* expression and greater resistance to apoptosis [47].

**Fig. 6.**
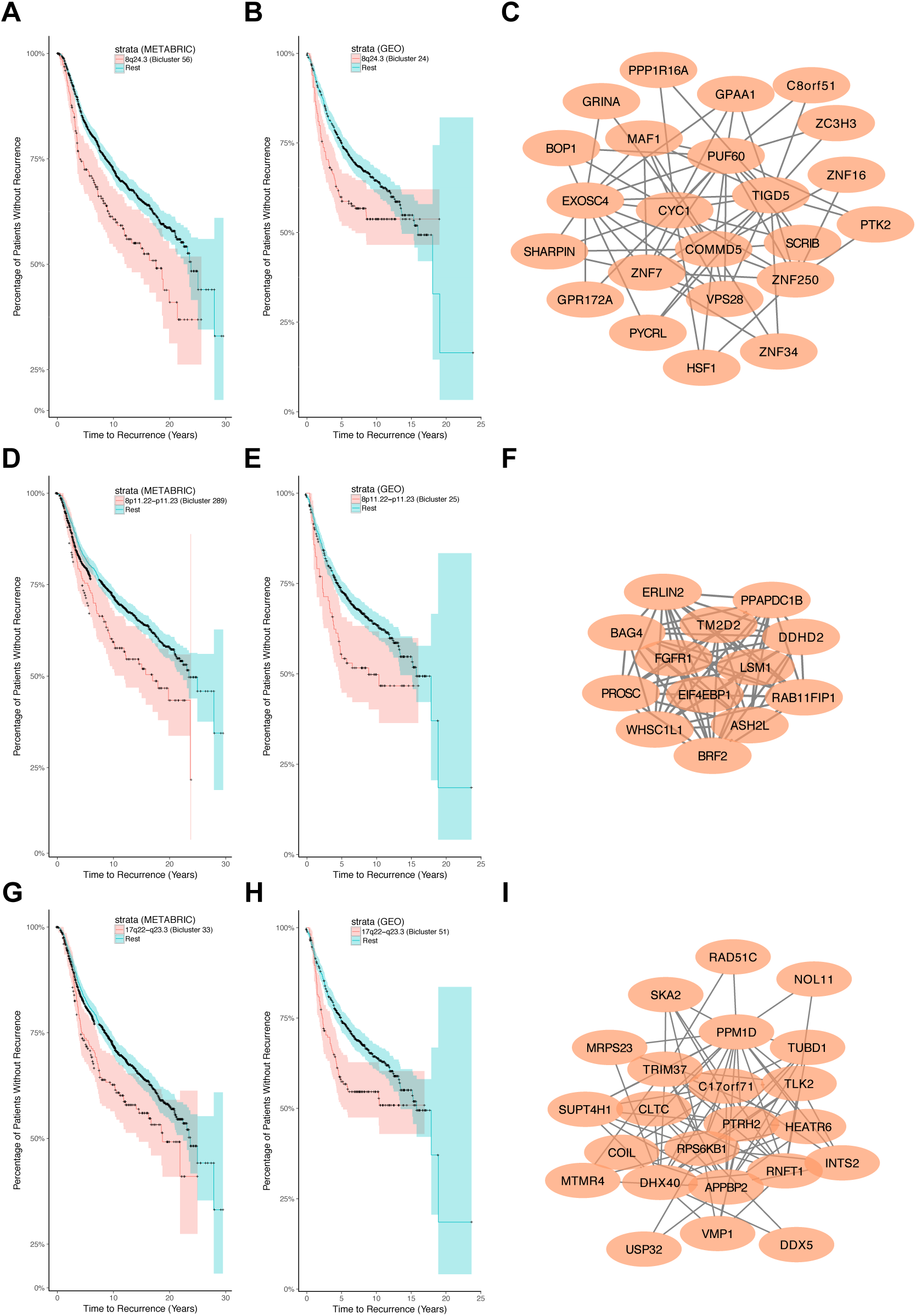
Clinically pertinent biclusters. Kaplan-Meier survival curves for the set of patients in the bicluster (red) compared to the remaining set of patients (blue) for METABRIC and GEO datasets, together with the graphs corresponding to the biclusters for 8q24.3 (A, B, and C), 8p11.22-p11.23 (D, E, and F) and 17q22-q23.3 (G, H, and I).

For both METABRIC and GEO, patients in biclusters associated with copy number gains of the 8p11.21-p11.23 loci (METABRIC – bicluster 289, GEO – bicluster 25) had significantly shorter RFS times compared to patients without amplification of this locus (**Fig. 6D**, **6E** and **6F**). We found that patients in this bicluster were enriched in the Luminal B subtype, which has poorer prognosis than the Luminal A subtype among ER+/HER2-tumors [48]. This suggested that amplification of the 8p11.21-p11.23 loci may be another marker of potential failure of ER targeted therapy.

Similarly, we found that patients whose tumors have copy number gains of the 17q22-q23.3 locus (METABRIC – biclusters 33 and 119, GEO – biclusters 15 and 160) had significantly shorter RFS times compared to patients whose tumors do not exhibit such a copy number gain (**Fig. 6G**, **6H** and **6I**). For METABRIC, this cohort was enriched in the Luminal B (PAM50), ER+/HER2+, and ER-/HER2+ subtypes (FDR < 0.001). For GEO, this cohort was enriched in the ER+/HER2+ and ER-/HER2+ subtypes (FDR < 0.05). This suggests that amplification of this locus may confer additional risk of recurrence in HER2+ breast cancers.

Note that the biclusters discussed above were not the only ones that exhibited differential relapse outcomes. For METABRIC, 61 biclusters out of 340 were found to exhibit differential relapse outcomes for the patients present in the biclusters. Out of these 61 biclusters, 69% were enriched in the ER-/HER2-subtype (64% for basal-like) with a significant proportion (67%) of these associated with copy number gains. For GEO, there were 48 such biclusters (13%) that exhibited differential relapse outcomes, 25% of these were enriched in the ER-/HER2-subtype.

Tests for enrichment of biclusters in tumors of higher grades revealed that 8 biclusters from TCGA were enriched in tumors of grade 3C. Some of these biclusters were associated with GO-BP terms related to angiogenesis, vasculogenesis, blood vessel maturation etc. For METABRIC, 4 biclusters were enriched in tumors of grade 3, out of which 2 were associated with the HER2 amplicon (17q12). For GEO, 68 biclusters were enriched in tumors of grade 3, including biclusters associated with CNA at the HER2 amplicon.

We also looked at the lymph node status of patients and observed that 4 biclusters in TCGA were enriched in samples with positive lymph node status in the corresponding patients. One was associated with the HER2 amplicon, while the others were associated with CNA at the 8q22.1-q22.3 loci, 17q23.1-q23.3 loci and the 19q13.43 locus, respectively. Similarly in METABRIC, we observed 4 biclusters enriched in samples with positive lymph node status in the corresponding patients - 2 of them were associated with copy number gains at the HER2 amplicon, the other 2 were associated with copy number gains at 19q13.11-q13.12 and 1q21.3-q25.1, respectively. Interestingly, biclusters associated with CNA at 8q24.3, 8p11.21-p11.23, and 17q22-q23.3 that exhibited poor RFS outcomes were not enriched in tumors of higher grades or in patients with positive lymph node status in any of the 3 datasets. In case of METABRIC, we additionally confirmed that none of these biclusters (8q24.3, 8p11.21-p11.23, 17q23.1-q23.3) were among the 36 biclusters enriched in samples with the poorest expected 5-year survival outcome (Nottingham Prognostic Index (NPI): > 5.4) [49, 50]. This highlights the importance of these altered transcriptomic signatures for reclassification of patients into the category with higher risk of recurrence.

### Hierarchical clustering of biclusters reveals shared mechanisms

Sample membership based hierarchical clustering of biclusters revealed distinct groups of biclusters that presumably share common functional mechanisms (**Fig. 7**). These included clusters associated with cell cycle and proliferation, immune response, cell adhesion (extracellular matrix), translation, mitochondrial translation, and ribosomal RNA processing pathways. Since a significant fraction of our biclusters were associated with copy number alterations, we also found distinct groups of biclusters associated with significant copy number changes such as the ones associated with the HER2 amplicon, the 8p11.21-p11.23 loci, or the 8q24.3 locus.

**Fig. 7.**
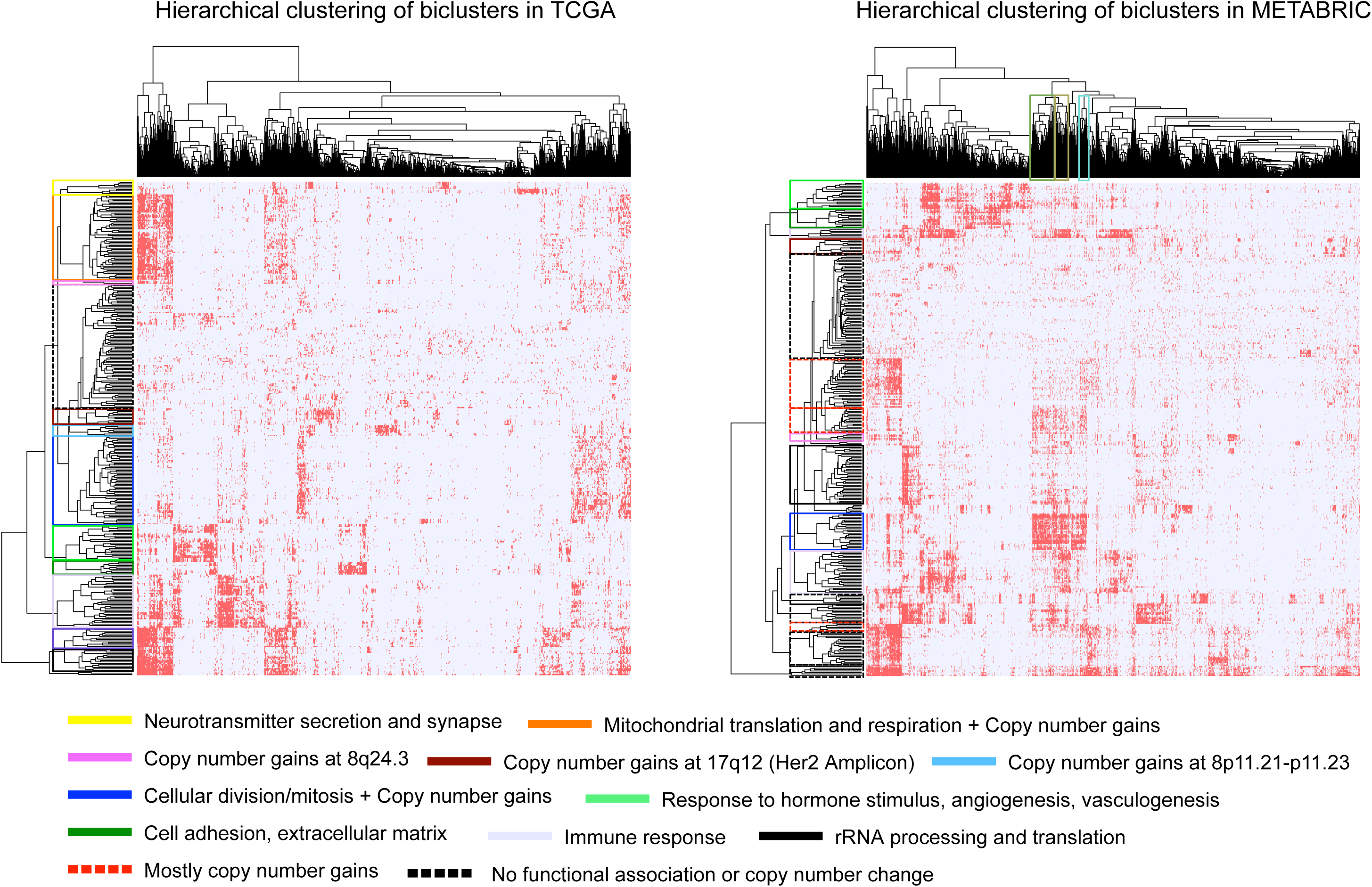
Hierarchical clustering of biclusters-samples. Hamming distance was applied to the biclusters-samples binary matrix for (A) TCGA and (B) METABRIC RFS datasets, respectively. The clusters of samples marked by green, brown, and cyan on top in panel (B) exhibit poor recurrence free survival. The green and brown clusters are associated with copy number gains at 8q24.3, while the cyan cluster is associated with copy number gains at 17q25.1-q25.3. Additionally, all three of them were enriched in gene signatures associated with cellular division and proliferation.

Similarly, we used hierarchical clustering to group samples that were enriched in similar sets of biclusters, highlighting differential clinical outcomes. In particular, we observed 2 sets of samples enriched in biclusters associated with CNA at the 8q24.3 locus. In one group, the samples were enriched in biclusters related to immune response; this group showed significantly lower incidence of recurrence compared to those without enrichment in immune response-related biclusters. Both of these sets of samples were enriched in biclusters associated with cell division and proliferation. In contrast, we observed a cluster of samples enriched in biclusters associated with 8q24.3 copy number gain and a number of other loci, however these were not enriched in biclusters associated with cell division and proliferation. This group exhibited low incidence of recurrence. We also observed a cluster of samples with significantly poor RFS that were enriched in biclusters associated with CNA at 17q25.1-q25.3, and in biclusters associated with cell division and proliferation.

### TuBA compared to other biclustering methods

TuBA’s proximity measure distinguishes its biclusters from those identified by other algorithms by leveraging the size of the datasets to identify subsets of tumor samples that co-express subsets of genes at their most extreme levels (high or low) relative to other samples. We emphasize that TuBA is designed to identify biclusters with samples that correspond to the extremals for the corresponding sets of genes, and does not consider other subset of conditions for the same sets of genes for biclustering. In contrast, most biclustering methods seek sub-matrices with constant, or coherent gene expression patterns. Given this key difference, only those biclusters that exhibit such expression patterns in the extremal (top or bottom) subsets of samples for some subsets of genes, are expected to have agreement with the biclusters identified by TuBA. Therefore, a direct comparison between the biclusters discovered by other algorithms, with the ones identified by TuBA would necessarily be limited.

In their paper on DeBi [51], a novel biclustering method that identifies differentially expressed biclusters based on a frequent itemset approach, the authors applied their method to both synthetic and real gene expression datasets, including diffuse large B-cell lymphoma (DLBCL) data comprising 661 genes and 180 samples [52]. Apart from DeBi, they applied ISA [53], OPSM [54], QUBIC [55], and SAMBA [56] to this dataset; we used these results to evaluate TuBA against these methods. In order to ensure a uniform and unbiased comparison between the enrichment results for biclusters from different algorithms, we used GeneSCF [35] to perform GO-BP enrichment on the biclusters obtained by all methods. **Fig. 8A** shows the proportions of GO-BP-enriched biclusters for five different significance levels (FDRs) – 0.001%, 0.1%, 0.5%, 1%, and 5%. For the FDR cutoff of less than 5%, almost all the biclusters for every algorithm were enriched in at least one GO-BP term. TuBA had 2 non-enriched biclusters out of 94, SAMBA had 2 non-enriched biclusters out of 128, and QUBIC had 1 bicluster out of 100 that were not enriched in a GO-BP term (**Supplementary Table 7**). As the FDR cutoff was lowered, TuBA had lower proportions of enriched biclusters compared to other algorithms for the corresponding FDR cutoffs. This can be partly attributed to the fact that the other algorithms discover biclusters that can have arbitrary overlaps between their genes. Since most of biclusters discovered by other algorithms shared genes with other biclusters, we could expect a certain amount of redundancy in enriched GO-BP terms. In contrast, TuBA precludes any overlap between the genes of the seeds of its biclusters. Moreover, its biclusters often include proximally located genes with aberrant expression due to copy-number changes, which may not show enrichment in GO-BP terms.

**Fig. 8.**
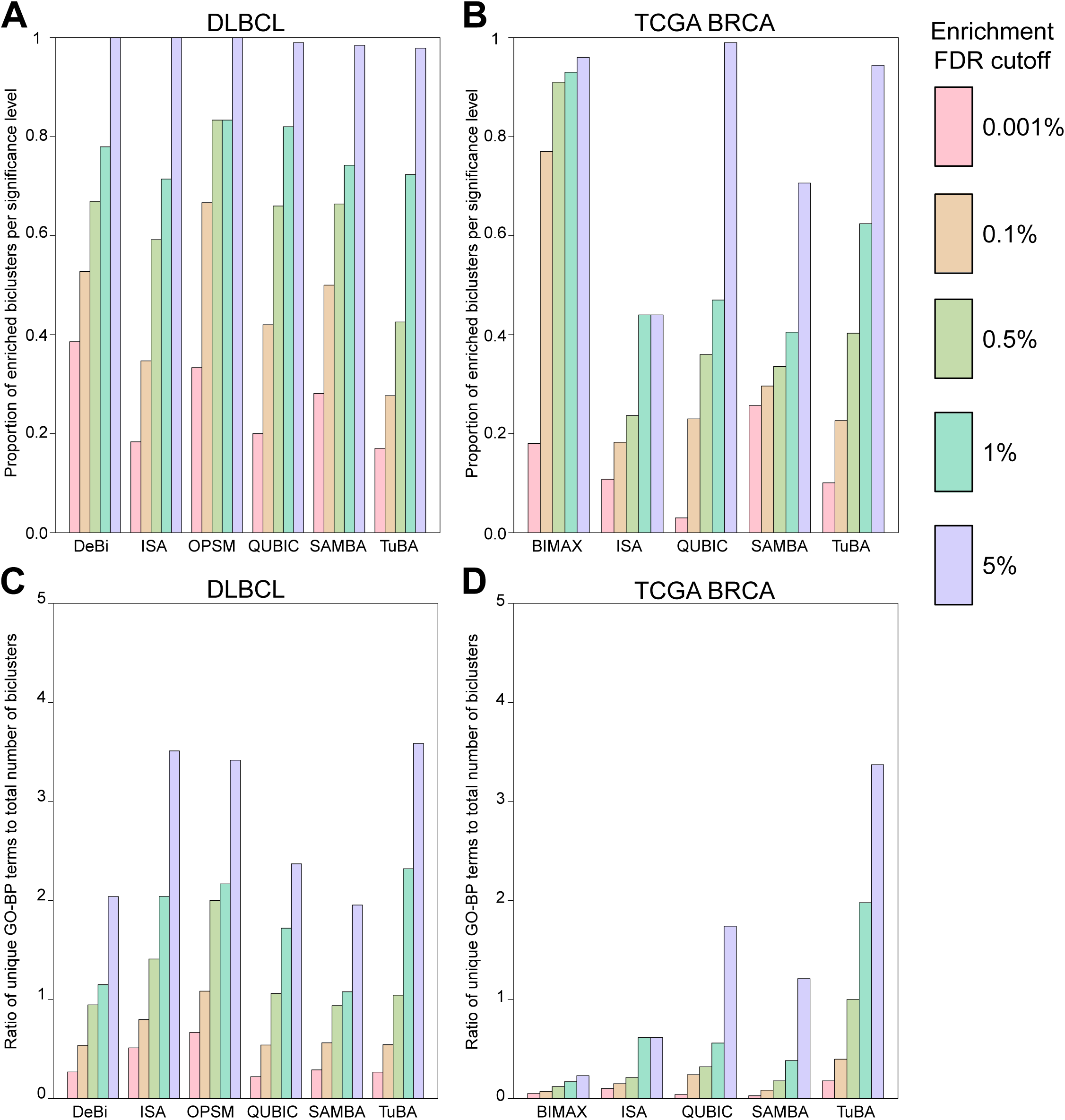
TuBA compared to other biclustering methods. Proportions of GO-BP terms enriched biclusters for each biclustering method at five different significance levels for (A) the DLBCL dataset, (B) the TCGA BRCA dataset. Ratios of number of unique GO-BP terms and total number of biclusters at five different significance levels for the (C) DLBCL dataset, (D) the TCGA BRCA dataset.

For a closer examination of the redundancy in the GO terms enrichment, we identified the top-five GO-BP terms for every bicluster obtained by each algorithm (not every bicluster was enriched in five distinct GO-BP terms, some had less than five, while others were not enriched in any term). For each algorithm, we prepared lists of all the unique GO-BP terms for the entire set of biclusters. The ratios of the number of elements in these lists to the total number of biclusters for each algorithm at five different significance levels show that TuBA identified biclusters enriched in a more extensive array of biological process terms (**Fig. 8B**).

In addition to the DLBCL dataset, we also analyzed the TCGA dataset with the following biclustering algorithms: (i) BIMAX [57], (ii) ISA, (iii) QUBIC, and (iv) SAMBA, using their respective default parameters (**Supplementary Tables 7 and 8**). For succinct descriptions of each of these algorithms we refer the reader to Prelic et al [8] and Pontes et al [55]. We used the biclust package in R for BIMAX [58], the isa2 package in R for ISA [59], the QUBIC package in R for QUBIC [60], and the Expander software for running SAMBA [61]. **Fig. 8C** shows the proportion of GO-BP terms enriched in biclusters of each algorithm for five different significance levels. TuBA compared favorably with other algorithms, especially when we accounted for the redundancy of the GO terms that were found enriched in the biclusters. We observed again that TuBA’s biclusters were enriched in a larger set of distinct biological process terms (**Fig. 8D**). We observed similar results for METABRIC (**Fig. S8A**, **S8B**). The results of the comparative analysis are summarized in **Supplementary Methods**.

In these analyses, the choice of the parameters is a crucial factor in determining the performance of each biclustering method. It is possible that different results could be obtained by more prudent choices of parameters for the other algorithms, however a detailed analysis of optimal parameter choices for each of these algorithms is beyond the scope of this study. We must point out for TuBA, that for any given dataset, there is no optimal (or default) choice of its two parameters; the biclusters obtained for any given choice of the parameters simply satisfy the basic requirements laid down by those choices. We looked at GO-BP-term enrichments for TuBA’s biclusters for five different choices of the overlap cutoff (**Supplementary Methods**). Although, the total number of biclusters obtained differed for each choice, the proportion of enriched biclusters at different significance levels remained similar irrespective of the parameter choice (**Fig. S9A**). Similarly, the ratio of the number of unique GO-BP terms and the total number of biclusters was consistent across all five choices of overlap cutoffs (**Fig. S9B)**. In earlier studies comparing biclustering algorithm [8, 57], synthetic datasets were generated with constant, shifting, and/or scaling patterns of expression for subsets of conditions and genes. The algorithms were evaluated based on how well they were able to identify the known biclusters implanted in these synthetic datasets. Since TuBA is not based on a mathematical model of the data in its biclusters, a comparison based on synthetic datasets is not feasible. However, in case of tumor datasets we have the benefit of complementary genomic data that could provide us with truth-known scenarios for validation. For example, alterations at the genomic level can directly influence the expression levels of genes; it is well known that a significant proportion of tumors across multiple tumor types frequently exhibit genomic alterations such as gains or losses in the copy numbers of genes. Quite often, these alterations are not limited to a single gene but include multiple genes located at neighboring chromosomal locations. If such alterations are located at transcriptionally active sites, then co-expression of the neighboring genes that are affected by it will be observed. In BRCA for instance, approximately 15-20% of tumors possess extensive gains in copy numbers of genes at the 17q12 cytoband locus (includes *ERBB2 (Her2)*, *STARD3*, *GRB7*, *PNMT, PGAP3, MED1* etc.). Identification of co-expression of genes at this locus in the subset of samples that are histologically HER2-positive (HER2+) represents a simple truth-known scenario that can be used to verify whether a given biclustering algorithm identifies the co-expression of these genes in the subset of samples that exhibit this alteration. We identified HER2+ samples in the TCGA dataset, and for each biclustering algorithm selected those biclusters that were enriched in these samples (hypergeometric test FDR < 0.001). BIMAX and SAMBA did not discover any, but ISA identified two biclusters (biclusters 71 and 72) enriched in HER2+ samples. Although the genes from the 17q12 amplicon – *ERBB2, STARD3, GRB7, PNMT, PGAP3, MED1* etc. were present in ISA’s enriched biclusters, they comprised a small subset within the genes in them – bicluster 71 had 639 genes, while bicluster 72 had 539 genes, respectively. QUBIC also had four biclusters that were enriched in HER2+ samples, however they did not contain any genes from the HER2 amplicon (including *ERBB2*). In contrast, not only did TuBA identify a bicluster (bicluster 256) exclusively associated with the HER2 amplicon, it identified many other biclusters associated exclusively with CNA of genes located near each other.

In summary, apart from TuBA, only ISA identified co-expression of the genes located at the HER2 amplicon. However, ISA’s co-expression module corresponding to the amplicon was embedded within much larger sets of genes. In the absence of information about copy number gain of the *ERBB2* gene, it would be a challenge to explicitly identify the co-expression module corresponding to the amplicon, and in turn infer the underlying mechanism for their co-expression. TuBA successfully uncovers those co-expressed sets of genes that are associated with CNA of neighboring sites on the chromosome, and is particularly efficient at identifying transcriptionally active copy number gains, as compared to other algorithms.

The nature of our proximity measure allows us to determine differential co-expression signatures without the need to specify subsets of samples in advance. Gao *et. al.* [62] proposed a biclustering method, *Bicmix*, based on a Bayesian statistical model to infer subsets of co-regulated genes that covary in all samples, or in only a subset of samples. They also developed a principled method to recover context-specific gene co-expression networks from the sparse biclustering matrices obtained by Bicmix. They applied Bicmix to the breastCancerNKI dataset and identified 432 genes that were differentially co-expressed in ER+, and ER-samples. Out of these 432 genes, 430 were up-regulated in ER-samples and down-regulated in ER+ samples, while 2 genes are down-regulated in ER-samples and up-regulated in ER+ samples. We applied TuBA (for high expression) to the same dataset with the following choice of parameters: (i) Percentile set size: 10%, and (ii) Overlap significance cutoff: FDR ≤ 1e-08. We obtained 549 biclusters, several of which comprised solely of probes that correspond to the same gene (**Supplementary Table 5**). This is reasonable, since probes corresponding to the same gene are expected to demonstrate higher expression levels in the same set of samples. We inquired whether some of the biclusters discovered by TuBA corroborated the differential co-expression signature between ER+ and ER-samples identified by Bicmix. Using Fisher’s exact test, we determined that the set of 430 genes up-regulated in ER-samples and down-regulated in ER+ samples were enriched in 30 biclusters discovered by TuBA – bicluster 5 shows the maximum enrichment (FDR < 1e-165). In fact, the genes that had the highest degrees in the co-expression network discovered by Bicmix – *CD247*, *CD53*, *IL10RA*, and *CXCR3* – were among the ones with highest degrees in bicluster 5 discovered by TuBA. The two genes (*SFRP2* and *COL12A1*) that were up-regulated in ER+ samples and down regulated in ER-samples were also found to be co-expressed in a TuBA bicluster (bicluster 115). TuBA also identified biclusters corresponding to amplicons at 17q12 (HER2), enriched in ER-samples (FDR = 0.02); 8q24.3, enriched in ER-(FDR = 0.003) samples; 17q25-q25.3, enriched in ER-samples (FDR = 7.09e-05). Thus, in addition to the differential co-expression network identified by Bicmix, TuBA recovers biclusters associated with genomic alterations such as CNA, several of which are differentially expressed between ER+ and ER-samples. Overall, TuBA recovered 144 biclusters enriched in ER-samples, and 31 biclusters enriched in ER+ samples (FDR < 0.05). This is consistent with our earlier observation that a significant proportion of biclusters discovered independently in the TCGA, METABRIC, and GEO datasets, were enriched in the ER-/HER2-subtype.

### Runtime Analysis

TuBA’s graph-based algorithm relies on the identification of largest cliques, which is a computationally hard problem. Large graphs (both in terms of the number of genes, and edges) can potentially lead to long computation times. The size of our graphs is principally determined by the choice of the cutoff for the second parameter – the significance level of overlap between percentile sets. We varied the cutoffs for the TCGA and METABRIC datasets, respectively, such that the total number of edges in the resultant graphs ranged between 10,000 and 250,000.

For choices of overlap cutoffs consistent with our suggested heuristic, we recorded TuBA’s computation time to generate final biclusters for each dataset (**Fig. 9A**). The computation time for TCGA increased dramatically as the number of edges in the graphs went beyond 150,000 edges. In particular, while the computation time for a graph with 200,000 edges for METABRIC was approximately 70 minutes, the computation time for a graph of similar size for TCGA was approximately 42 hours. Thus, although METABRIC is the larger dataset with 24,368 genes and 1,970 samples compared to TCGA’s 20,241 genes and 908 samples, more iterations were required to identify all the largest cliques in the graphs for TCGA given its respective choices of parameters. We therefore conclude that TuBA’s computation time depends on the nature and complexity of the graphs themselves.

**Fig. 9.**
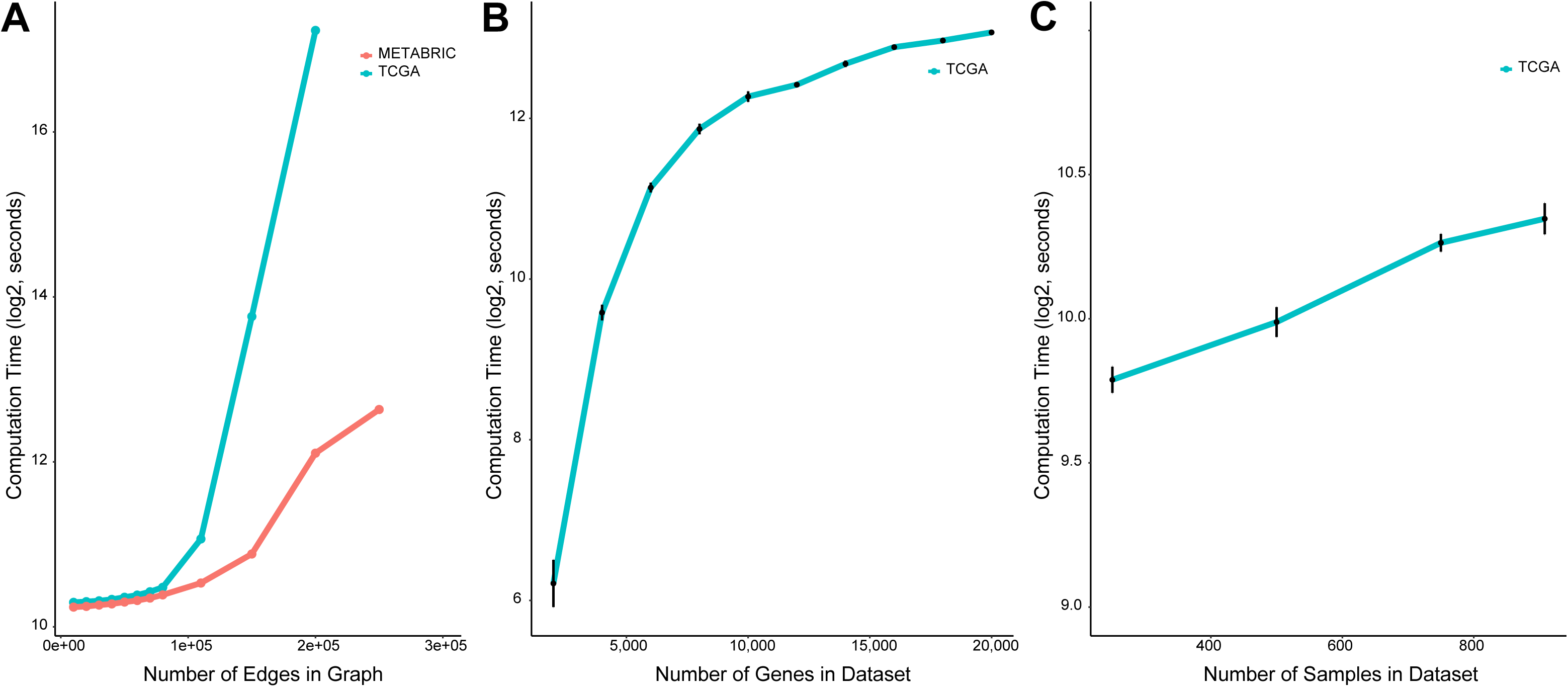
Runtime analysis of TuBA. (A) Computation time taken by TuBA to discover biclusters for different sizes (number of edges) of the graphs based on different choices of overlap cutoffs for the TCGA (blue), and METABRIC (red) datasets. (B) Dependence of computation time on the number of rows (genes) for choices of overlap cutoffs consistent with our suggested heuristic. Here, we chose overlap cutoffs consistent with our suggested heuristic, and ensured that comparable numbers of edges were generated for different datasets; this explains why the computation times for datasets with upwards of 14,000 genes were quite similar to each other. (C) Dependence of computation time on the number of columns (samples) for choices of overlap cutoffs consistent with our suggested heuristic.

We also investigated the impact of the size of datasets on computation time. We created new subsets from the TCGA dataset by randomly sampling a fixed number of genes. We varied the number of genes from 2,000 to 20,000, and created 10 randomly sampled datasets for each gene number (**Fig. 9B**). For each individual run, we chose an overlap cutoff consistent with our suggested heuristic, and ensured that comparable numbers of edges were generated for different datasets.

We also investigated the impact of the number of samples in a dataset. For this, we created five randomly selected subsets from the TCGA dataset each with 250, 500, 750, and 908 samples, (**Fig. 9C**). As expected, TuBA’s computation time did not depend strongly on the number of samples in the datasets.

In its current implementation, using a 2.7 GHz Intel Xeon processor, and 48 GB of RAM. TuBA has longer runtime than most other existing algorithms. Depending on the choice of the overlap cutoff, the runtimes can vary between 15 and 120 minutes for datasets with approximately 20,000 genes and 1,000 samples.

## DISCUSSION

Global clustering approaches have successfully unveiled distinct disease subtypes in tumors, prompting the community to look beyond traditional clinico-pathological signatures to identify relevant disease processes. However, the extensive heterogeneity, even within tumors of a given subtype, confounds the identification of many altered transcriptional programs by such unsupervised clustering methods.

In this paper, we introduce an algorithm called TuBA based on a proximity measure specifically designed to extract gene co-expression signatures that correspond to the extremes of expression (both high and low for RNASeq data, and high for array-based platform). This enables us to preferentially identify co-aberrant gene signatures associated with the disease states of tumors. The identification of altered transcriptional profiles can be particularly relevant for those tumors that have so far eluded targeted drug development for therapy. This is exemplified by tumors of the basal-like or triple negative subtypes for BRCA. Although these tumors account for only ~15% of all BRCAs in the population, a significant fraction of biclusters identified by TuBA corresponded to alterations associated with tumors of these subtypes. For each dataset, a simple estimation of enrichment of samples in a given bicluster within any other bicluster, revealed that the samples in the biclusters corresponding to CNA at 8p11.21-p11.23 or 17q12 were enriched (FDR < 0.001) independently in ~5% of all biclusters for both TCGA and METABRIC, respectively. In sharp contrast, 30–40% of all biclusters were enriched in samples with copy number gains at the 8q24.3 locus (FDR < 0.001). Additionally, 51% of all biclusters obtained from the low expression analysis of TCGA were enriched in the samples corresponding to the 8q24.3 bicluster. Previous studies have also identified the amplicon at 8q24.3 by Representational Difference Analysis as a location of oncogenic alterations in breast cancer that can occur independent of neighboring *MYC* amplifications [63]. Although the 8q24.3 bicluster itself is enriched with ER-/HER2-samples, these observations, together with poor RFS outcome observed independently in both METABRIC and GEO, highlight this locus as a promising prognostic marker for BRCA tumors, irrespective of subtype.

We must mention a notable exception in the biclusters discovered by TuBA for all three datasets – none of the biclusters contained the *ESR1* gene, which codes for estrogen receptor. Closer inspection revealed that *ESR1* had statistically significant associations with several genes, however the level of significance of overlap with these genes was much lower (FDR > 1e-07) than the chosen cut-off for all three datasets. While it is known that about 70% of BRCA exhibit elevated expression of estrogen receptor, over-expression of estrogen receptor may not be a sufficient condition to drive co-expression of genes involved in other pathways [64]. This may explain why co-expression of *ESR1* with other genes was not as significant as the other associations that were extracted and summarized in TuBA’s biclusters.

Apart from highlighting the heterogeneity of CNA-associated alterations in tumors of the ER-/HER2-subtype or basal-like subtype, TuBA offered a glimpse into the utility, the limitations, and the potential pitfalls with the current subtype classification approaches. In the ER/HER2-based subtype enrichment, we observed a significant proportion of biclusters that were not specifically enriched in any one of the four subtypes. For instance, several CNA-associated biclusters from chromosome eight were not subtype-enriched. In the case of PAM50 subtype classification however, we observed that most of these biclusters were enriched in the Luminal B subtype for METABRIC (and to a limited extent for TCGA). While this appears to indicate that PAM50 offers an improvement on the traditional clinico-pathological approach to subtype classification, it unfortunately fails to classify several samples associated with overexpression of ERBB2 as HER2-positive. As a consequence, several of our biclusters associated with the HER2 amplicon and copy number gains in the neighboring locations on chromosome 17 (17q.21.1-q21.2 and 17q21.32-q21.33), for both METABRIC and TCGA, were observed to be enriched in the Luminal B subtype. This corroborates the modest level of agreement with PAM50 classification reported in [37], as well as disagreements in later studies [65]. Given that trastuzumab is a clinically proven therapeutic drug for HER2+ tumors, misclassification of these patients into any other subtype can be highly disadvantageous.

Change in copy number is often not a sufficient condition for elevated (or suppressed) expression levels of transcripts, as there are multiple layers of regulation of transcription in cells [66, 67]. TuBA specifically identifies sets of genes with copy number changes that are transcriptionally active (or inactive), filtering out the ones that are unlikely to influence disease progression. Moreover, the graph-based approach allows us to infer the relative importance of each gene within a bicluster, based on its degree. In the case of high expression analysis, the degree of each gene is an indicator of how frequently it is expressed aberrantly at high levels by the subset of samples that comprise any given bicluster. As an example, consider the CNA-associated bicluster from TCGA corresponding to gains at the 8q22.1-q22.3 loci. The bicluster exhibited enrichment in lymph node positive patients (the corresponding bicluster in METABRIC has a significance level of *FDR* = 0.052 for patients with positive lymph node status). The gene with the highest degree in the bicluster was *MTDH* (metadherin), which has been shown to be associated with increased chemo resistance and metastasis in BRCA [68–70].

Clustering analysis of biclusters and samples based on the membership of samples within biclusters allowed us to identify the sites that were altered concomitantly within the same subsets of samples. Moreover, we improved our perspective on the tumor microenvironment in the subsets of samples that exhibit non-tumor associated signatures (such as immune, extracellular matrix, etc.). Differences in disease progression due to distinct microenvironments in tumors with similar transcriptional alterations can help us better understand the potential role of the microenvironment within the context of tumors harboring these specific alterations. For instance, we noticed a difference in RFS outcomes between two groups of patients that exhibit copy number gains at 8q24.3; the group that was additionally associated with an immune response signature was observed to have better RFS outcomes compared to the group that did not exhibit a strong association with the immune response.

Unlike most biclustering methods, TuBA does not allow arbitrary overlaps between its biclusters. This is because it is designed to discover biclusters with samples that correspond to the extremals for the corresponding gene set; biclusters with other conditions are not permitted for the same gene set. However, our biclusters are not exclusive, and some overlap between their genes and samples is permitted. For example, in case of an ER-/HER2-BRCA sample that exhibits CNA at 8q24.3, because of high immune-cell infiltration in the tumor, the same sample may also be present in the biclusters enriched in the sets of genes associated with immune response.

Another limitation of TuBA is that it can only be applied reliably for large datasets that contain at least 100 samples. Depending on cohort heterogeneity, some of the overlaps between percentile sets may not be significant in smaller datasets. However, the deliberate design of our proximity measure leveraging the size of the datasets offers a significant benefit – it not only enables the identification of the plethora of gene co-aberrations associated with the tumors, but also enables the estimation of the extent or prevalence of the identified alterations in the population. This is where the tunable aspect of TuBA becomes relevant – the two knobs should be viewed as valuable aids that help estimate the extents of the prevalence of various alterations in the tumor population and their clinical relevance. Although transcriptomic changes are not the ultimate determinants of progression, our algorithm holds the promise to improve therapeutic selection and design by identifying significantly altered transcriptional patterns associated with tumors.

## CONCLUSION

TuBA is quite distinct from other biclustering algorithms, in that, it is designed to identify biclusters with samples that correspond to the extremals for the corresponding sets of genes. Most biclustering algorithms are designed to identify nearly constant or coherent gene expression levels in subsets of genes across subsets of samples. However, we were able to show that TuBA performs outperforms other algorithms in identification of co-expressed genes located in transcriptionally active copy number altered sites. Moreover, from a differential co-expression perspective, TuBA offers an advantage over other methods, since no prior specification of subsets of samples (context) is necessary; the nature of our proximity measure ensures that such differential co-expression signatures are preferentially identified. Given these considerations, TuBA offers great promise as a biclustering method that can identify biologically relevant gene co-expression signatures that are not successfully captured by other unsupervised clustering or biclustering approaches. These signatures, along with the ones identified by other biclustering methods would enable a comprehensive understanding of the underlying alterations and shared mechanisms in subsets of tumors.

## Supporting information

Supplementary Tabl 1

Supplementary Tabl 2

Supplementary Tabl 3

Supplementary Tabl 4

Supplementary Tabl 5

Supplementary Tabl 7

Supplementary Tabl 8

## DECLARATIONS

### Ethics approval and consent to participate

Not applicable.

### Consent to publish

Not applicable.

### Availability of data and materials

TuBA is open-sourced and available in R scripts at https://github.com/KhiabanianLab. TCGA dataset was obtained from UCSC Xena Portal (http://xena.ucsc.edu). METABRIC dataset was obtained from the cBioPortal (http://www.cbioportal.org). GEO and breastCancerNKI datasets were obtained from Gyorffy & Schafer [24], and van’t Veer et al. [25] and van de Vijver et al. [26], respectively.

### Competing interests

The authors declare that they have no competing interests.

### Funding

HK acknowledges support from the American Cancer Society (IRG-15-168-01). The funding agencies had no role in the design of the study and collection, analysis, and interpretation of data or in writing of the manuscript.

### Authors’ Contributions

AS, GB, and HK conceived the study and designed the algorithm. AS implemented the algorithm and performed the statistical analyses. All authors contributed to the drafting of the manuscript and critical discussion of the results. All authors read and approved the final manuscript.

## Acknowledgements

This research was partially supported by the Biomedical Informatics Shared Resource at Rutgers Cancer Institute of New Jersey (P30CA072720) as well as Rutgers Office of Advanced Research Computing (NIH 1S10OD012346-01A1).

