## Supplementary Figures for "TuBA: Tunable Biclustering Algorithm Reveals Clinically Relevant Tumor Transcriptional Profiles in Breast Cancer"

Fig. S1

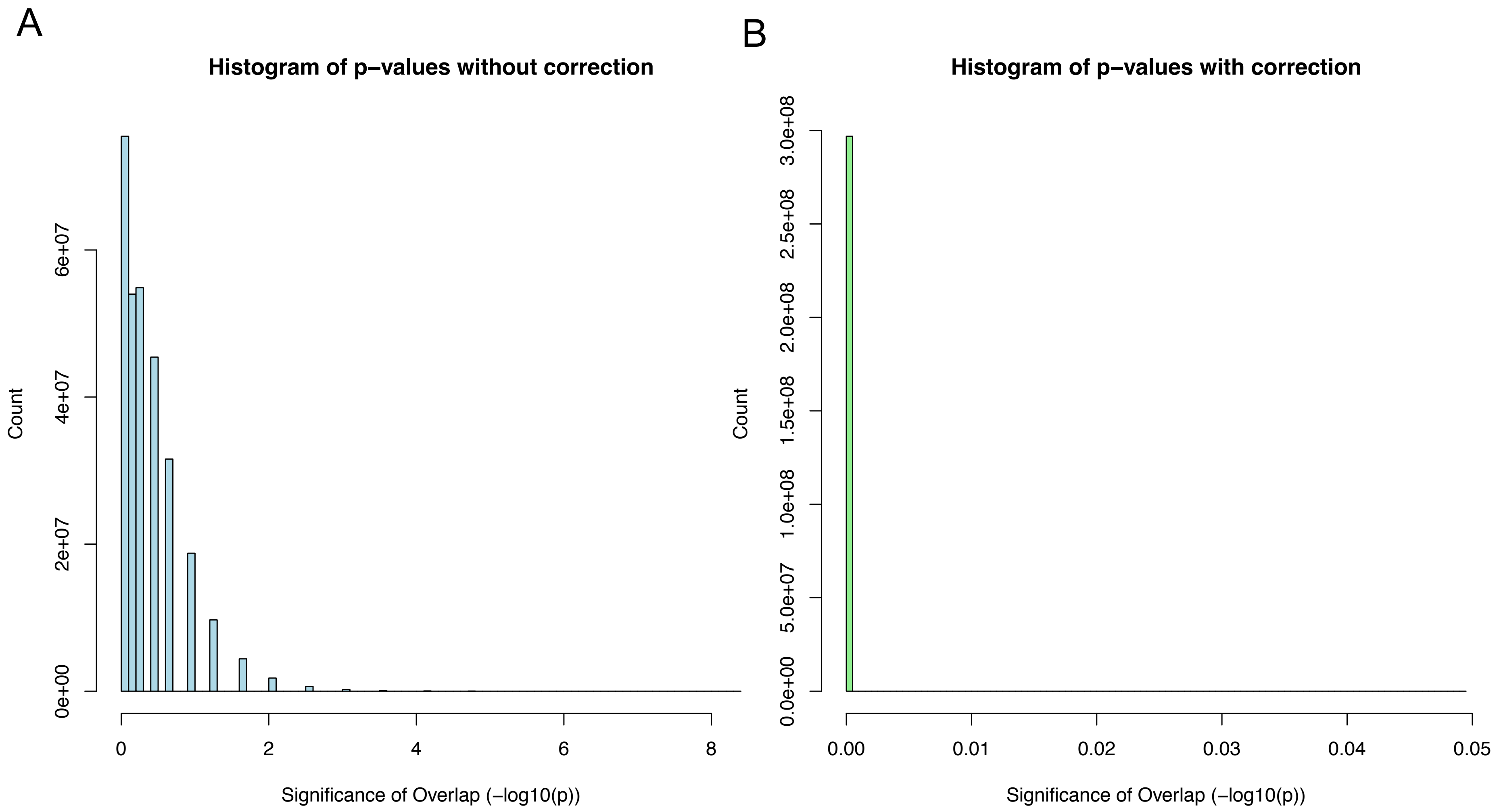
A
Histogram of p−values without correction
Count
0e+00
2e+07
4e+07
6e+07
0
2
4
6
8
Significance of Overlap (−log10(p))
B
Histogram of p−values with correction
Count
0.0e+00
5.0e+07
1.0e+08
1.5e+08
2.0e+08
2.5e+08
3.0e+08
0.00
0.01
0.02
0.03
0.04
0.05
Significance of Overlap (−log10(p))

Fig. S2

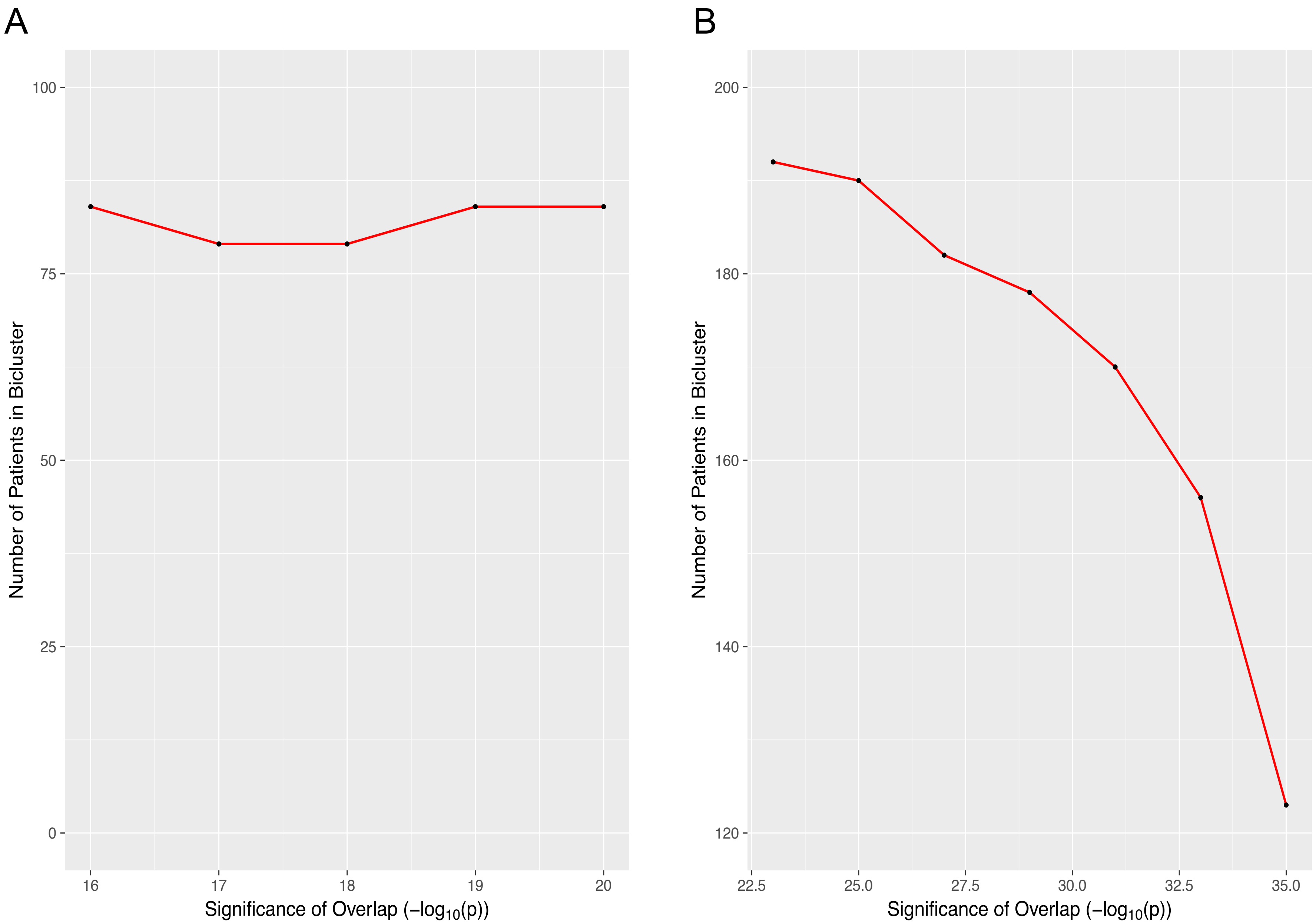
A
B
Number of Patients in Bicluster
100
75
50
25
0
16
17
18
19
20
Significance of Overlap (−log10(p))
200
180
160
140
120
22.5
25.0
27.5
30.0
32.5
35.0

Fig. S3

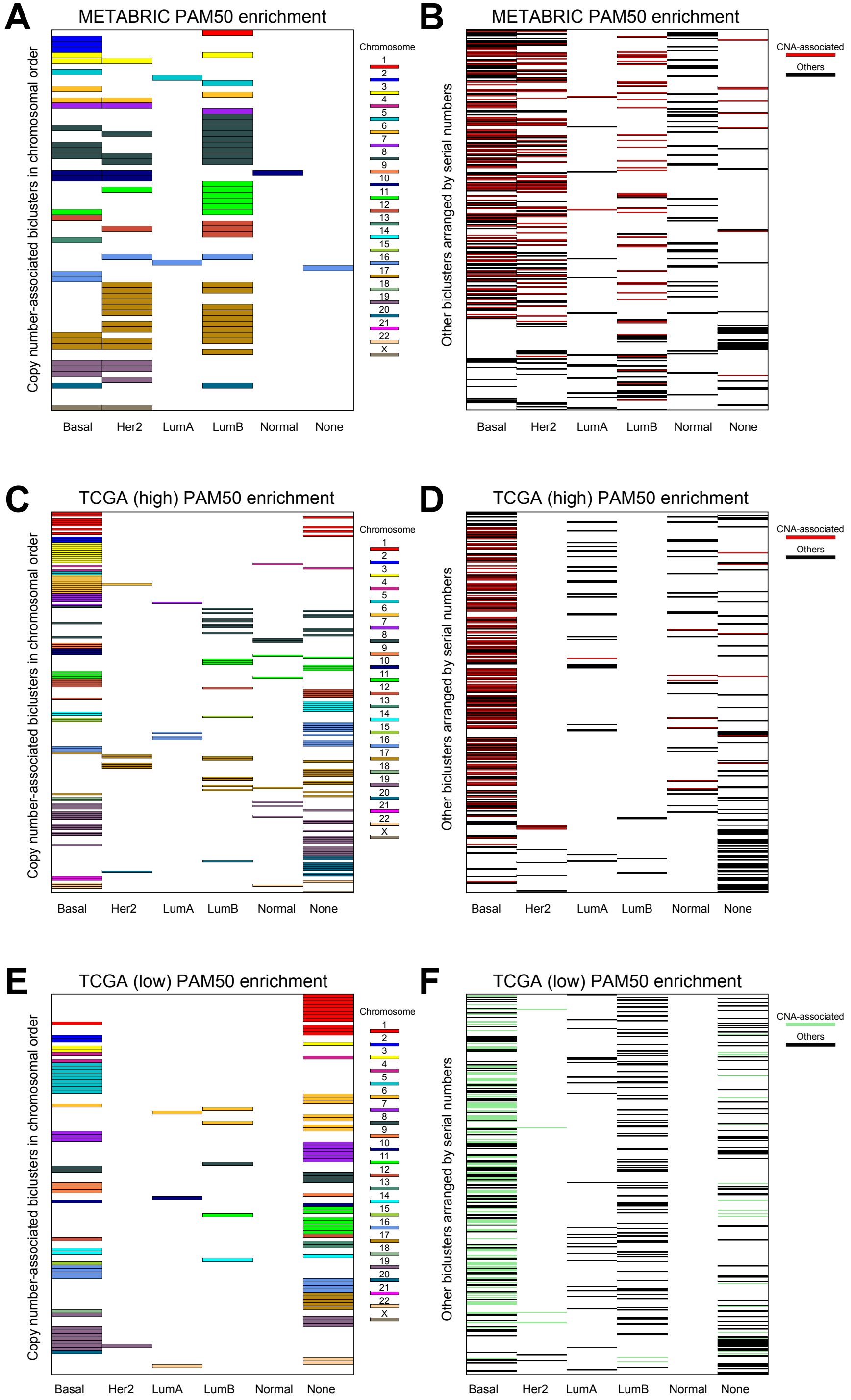
A
METABRIC PAM50 enrichment
Copy number-associated biclusters in chromosomal order
Chromosome
X
Basal
Her2
LumA
LumB
Normal
None
B
METABRIC PAM50 enrichment
Other biclusters arranged by serial numbers
CNA-associated
Others
C
TCGA (high) PAM50 enrichment
D
TCGA (high) PAM50 enrichment
E
TCGA (low) PAM50 enrichment
F
TCGA (low) PAM50 enrichment

Fig. S4

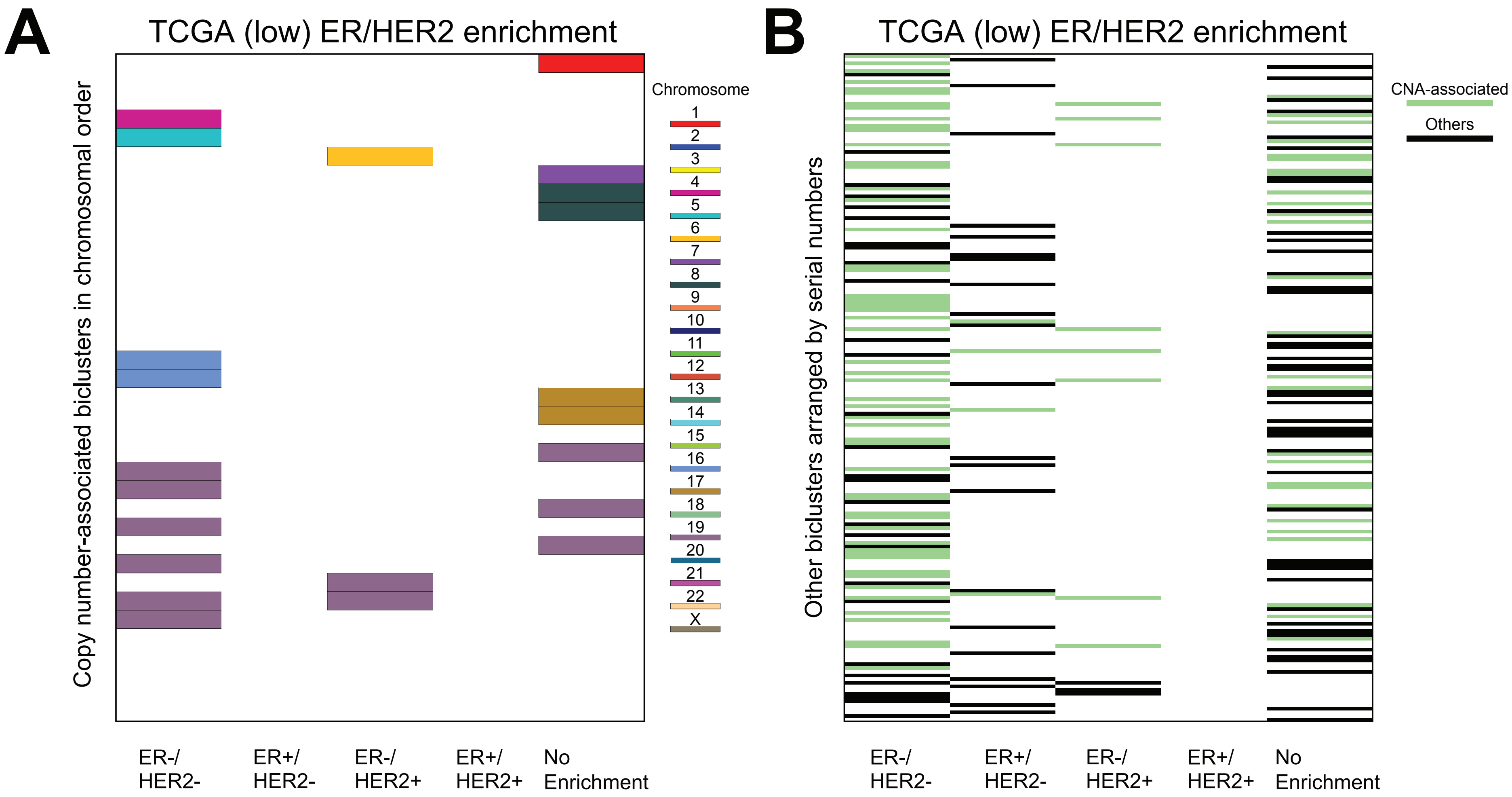
A
TCGA (low) ER/HER2 enrichment
Copy number-associated biclusters in chromosomal order
Chromosome
X
ER-/ HER2-
ER+/ HER2-
ER-/ HER2+
ER+/ HER2+
No Enrichment
B
TCGA (low) ER/HER2 enrichment
Other biclusters arranged by serial numbers
CNA-associated
Others
ER-/ HER2-
ER+/ HER2-
ER-/ HER2+
ER+/ HER2+
No Enrichment

Fig. S5

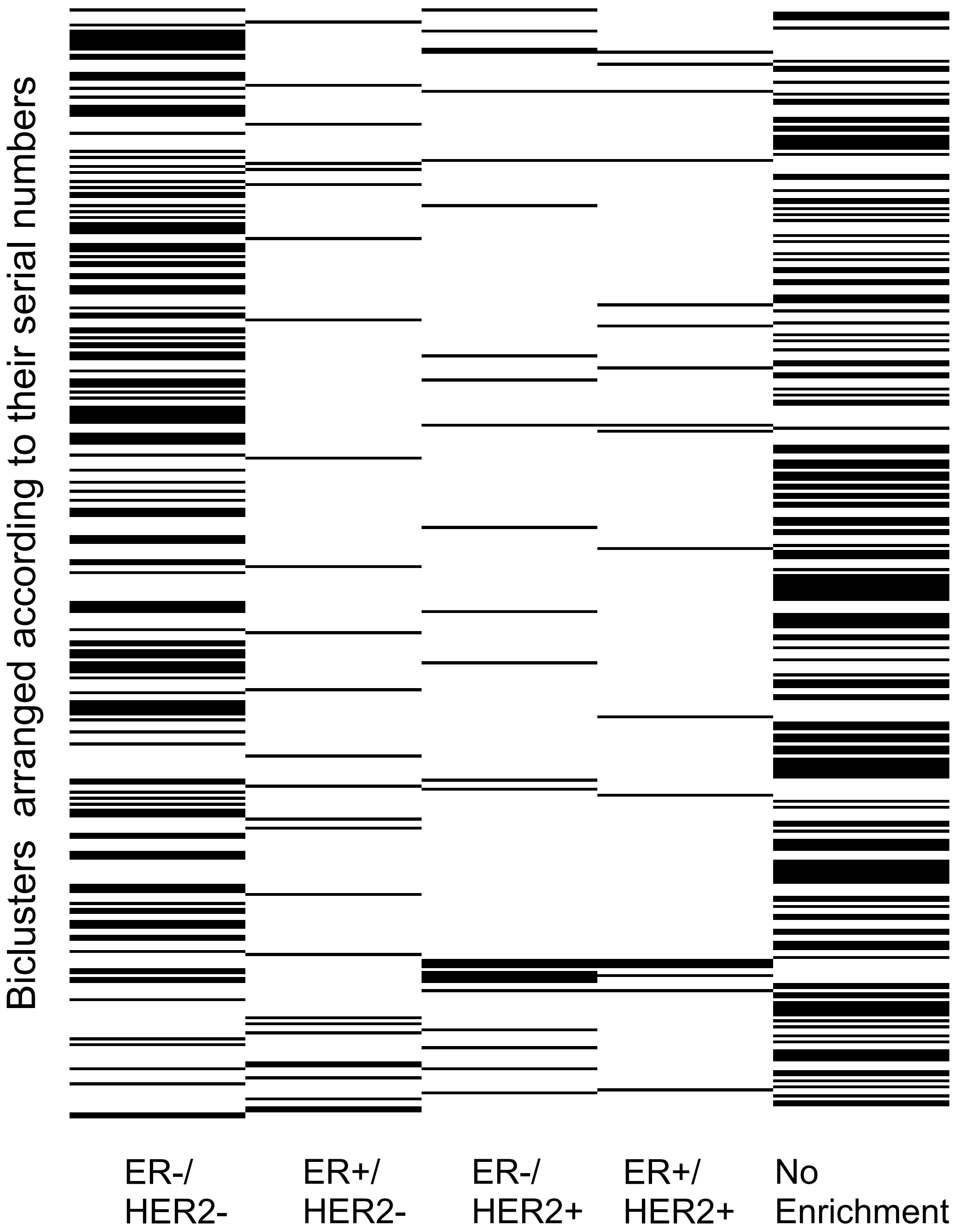
Biclusters arranged according to their serial numbers
ER-/ HER2-
ER+/ HER2-
ER-/ HER2+
ER+/ HER2+
No Enrichment

Fig. S6A

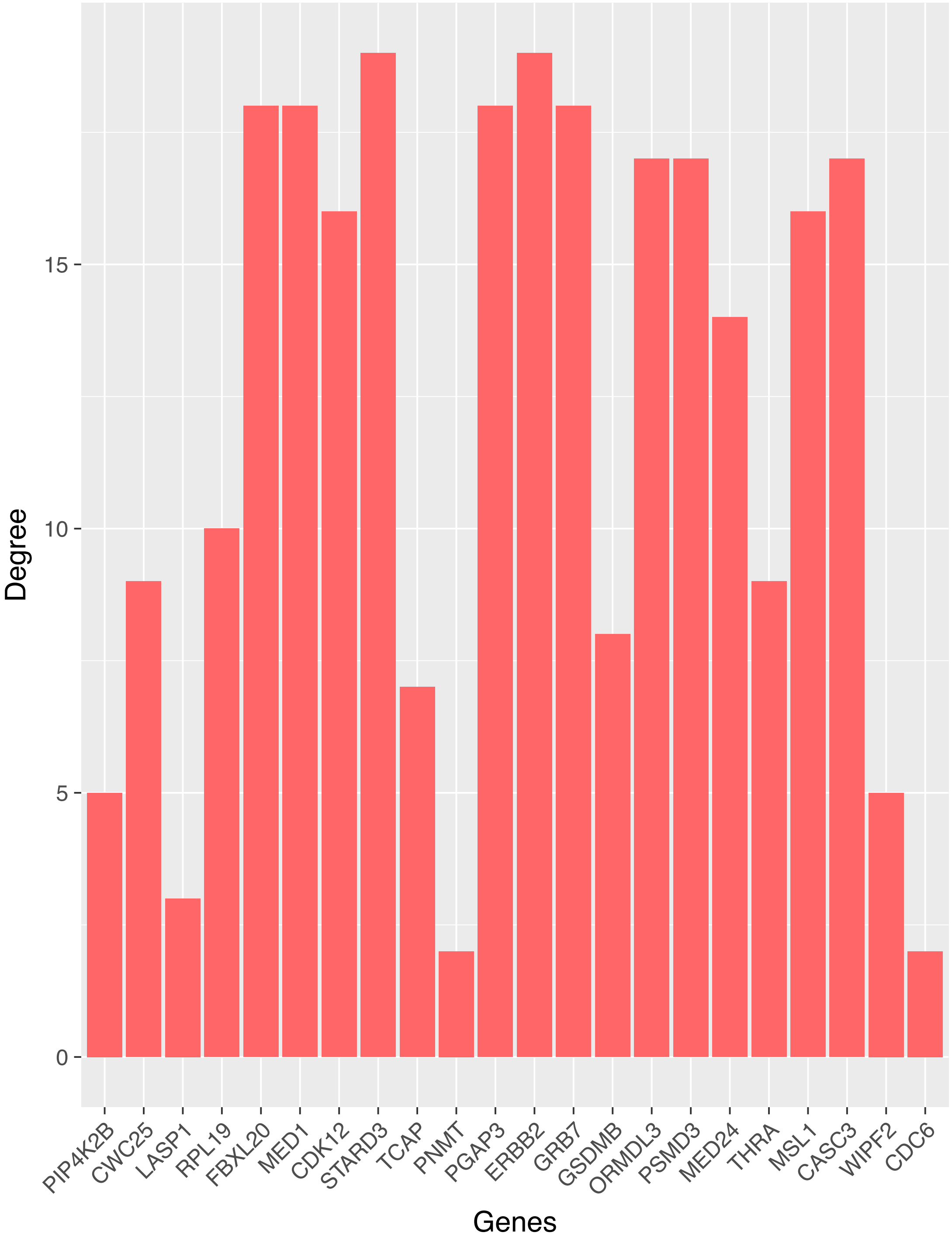
Degree
15
10
5
0
PIP4K2B
CWC25
LASP1
RPL19
FBXL20
MED1
CDK12
STARD3
TCAP
PNMT
PGAP3
ERBB2
GRB7
GSDMB
ORMDL3
PSMD3
MED24
THRA
MSL1
CASC3
WIPF2
CDC6
Genes

Fig. S6B

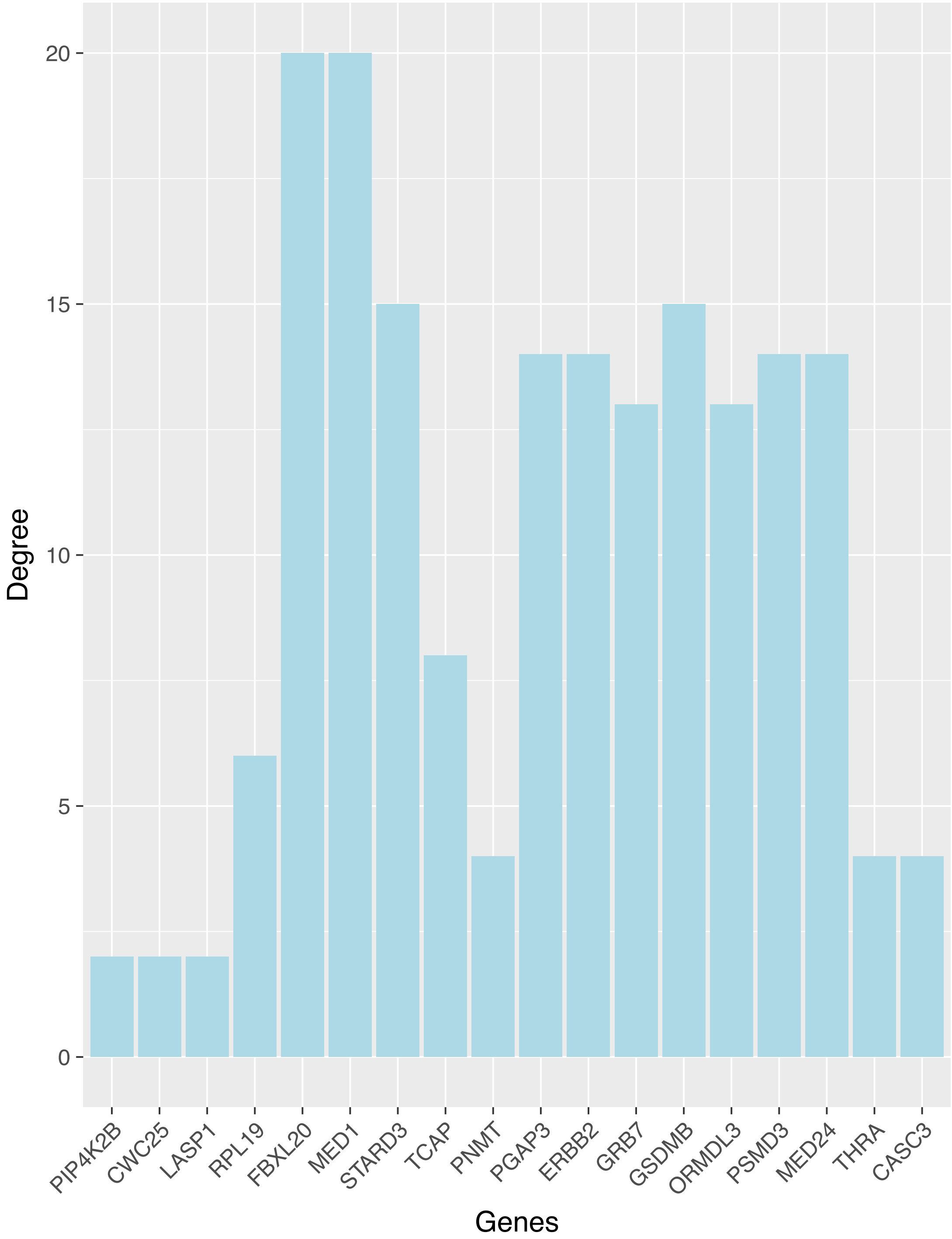
Degree
20
15
10
5
0
PIP4K2B
CWC25
LASP1
RPL19
FBXL20
MED1
STARD3
TCAP
PNMT
PGAP3
ERBB2
GRB7
GSDMB
ORMDL3
PSMD3
MED24
THRA
CASC3
Genes

Fig. S6C

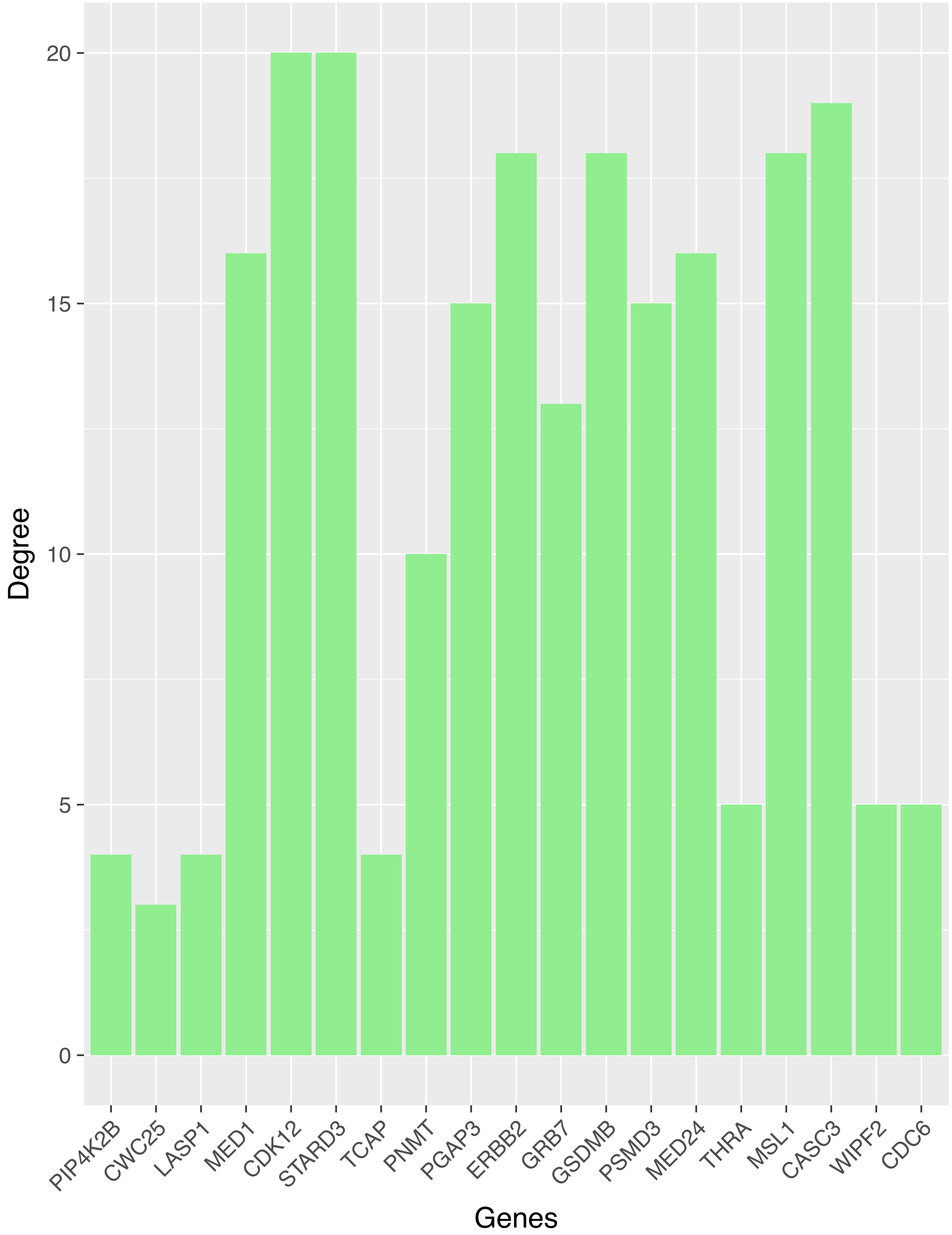
Degree
20
15
10
5
0
PIP4K2B
CWC25
LASP1
MED1
CDK12
STARD3
TCAP
PNMT
PGAP3
ERBB2
GRB7
GSDMB
PSMD3
MED24
THRA
MSL1
CASC3
WIPF2
CDC6
Genes

Fig. S7A

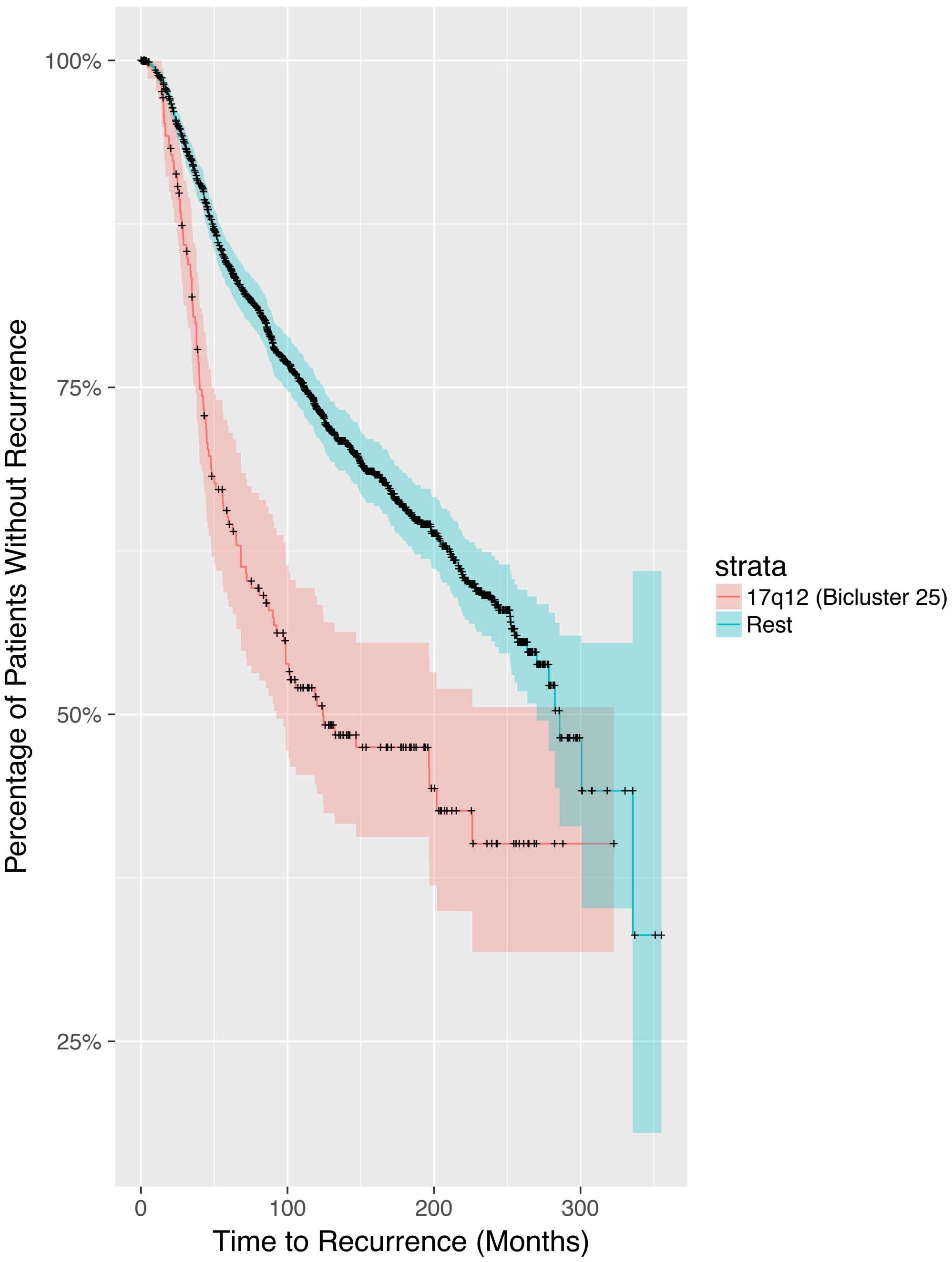

100%
75%
50%
25%
Percentage of Patients Without Recurrence
0
100
200
300
Time to Recurrence (Months)
strata
17q12 (Bicluster 25)
Rest

Fig. S7B

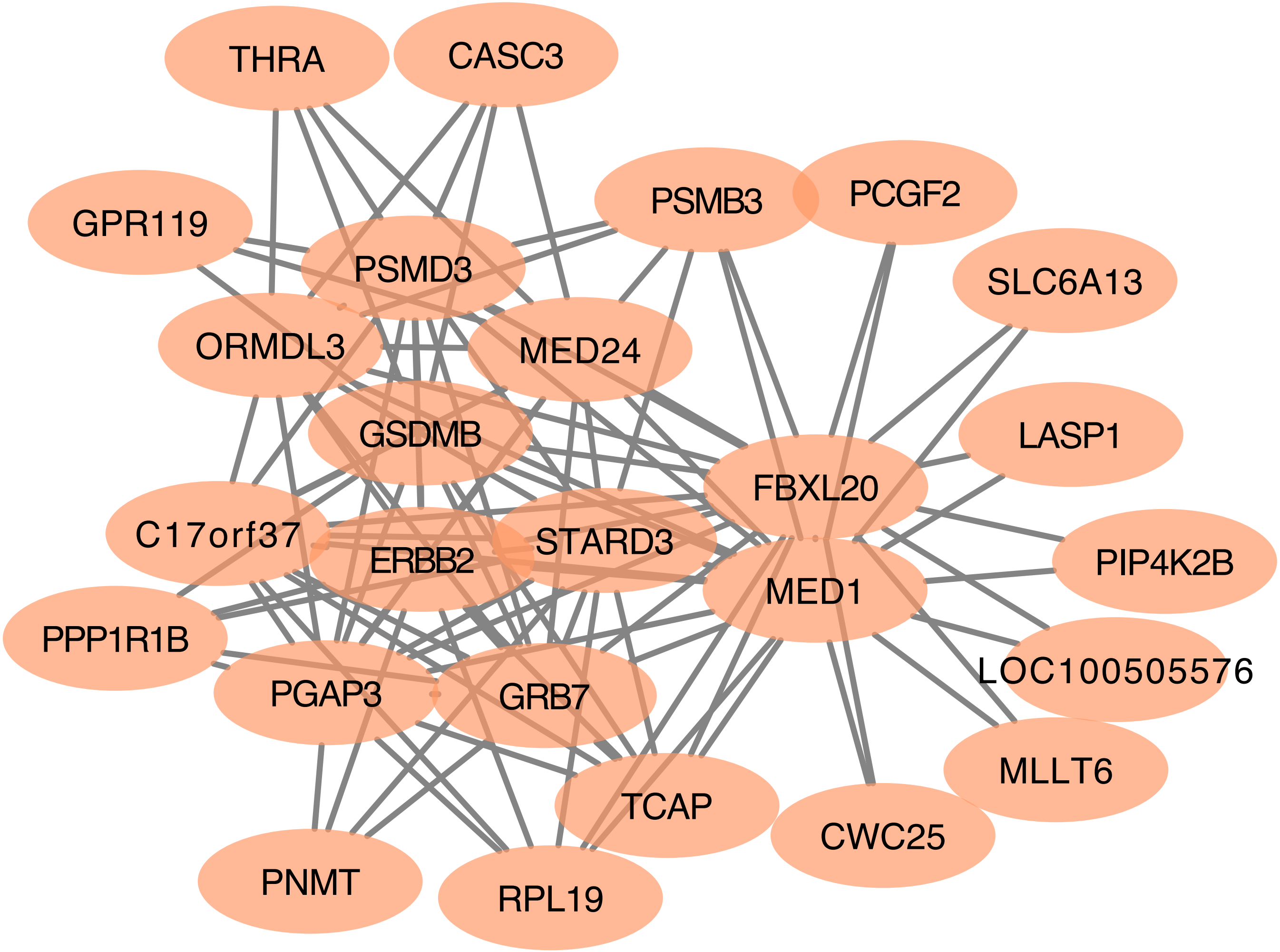
THRA
CASC3
GPR119
PSMB3
PCGF2
PSMD3
SLC6A13
ORMDL3
MED24
LASP1
GSDMB
FBXL20
C17orf37
STARD3
ERBB2
PIP4K2B
MED1
PPP1R1B
PGAP3
GRB7
LOC100505576
MLLT6
TCAP
CWC25
PNMT
RPL19

Fig. S8A

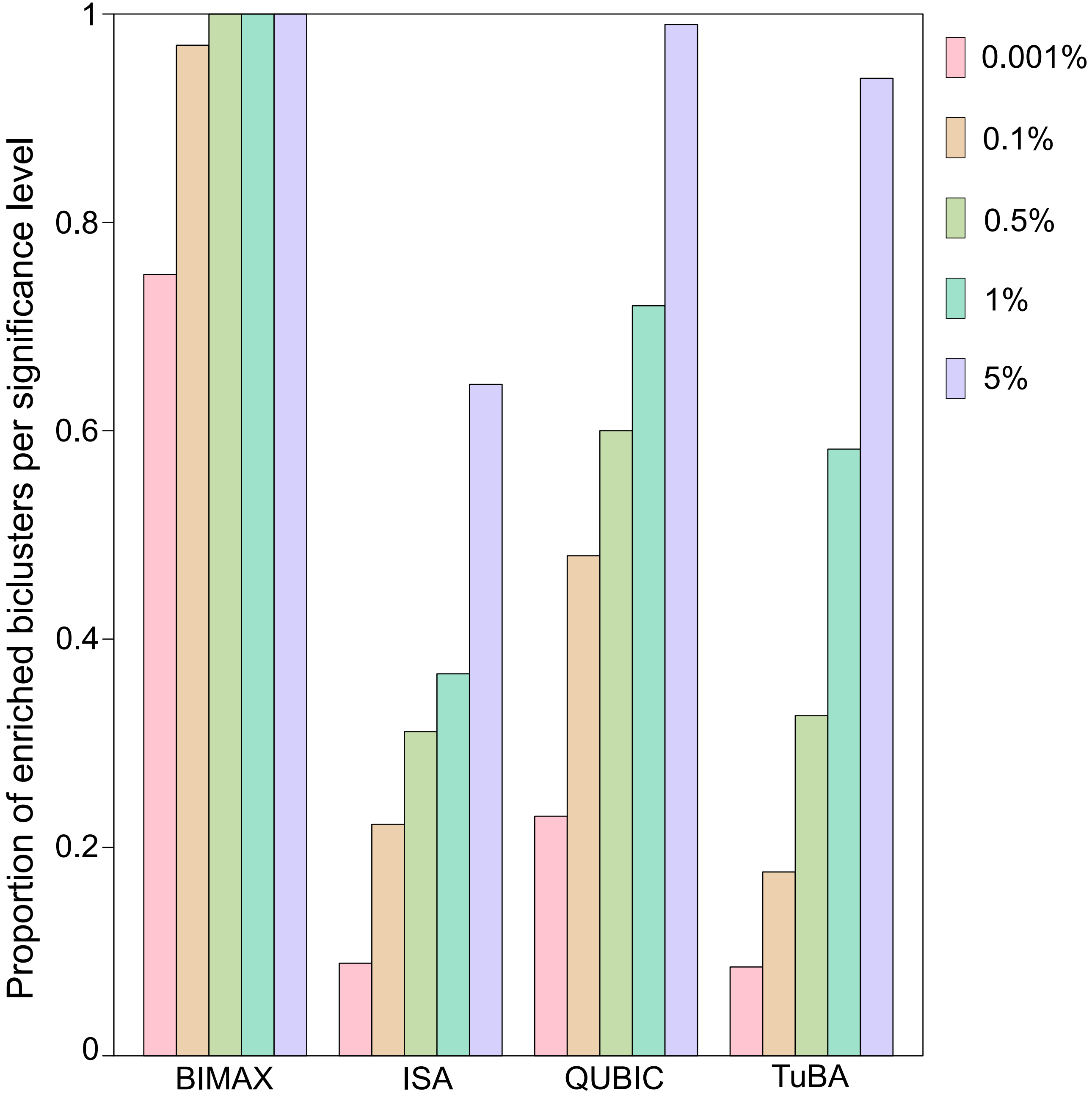

Proportion of enriched biclusters per significance level
0
0.2
0.4
0.6
0.8
1
BIMAX
ISA
QUBIC
TuBA
0.001%
0.1%
0.5%
1%
5%

Fig. S8B

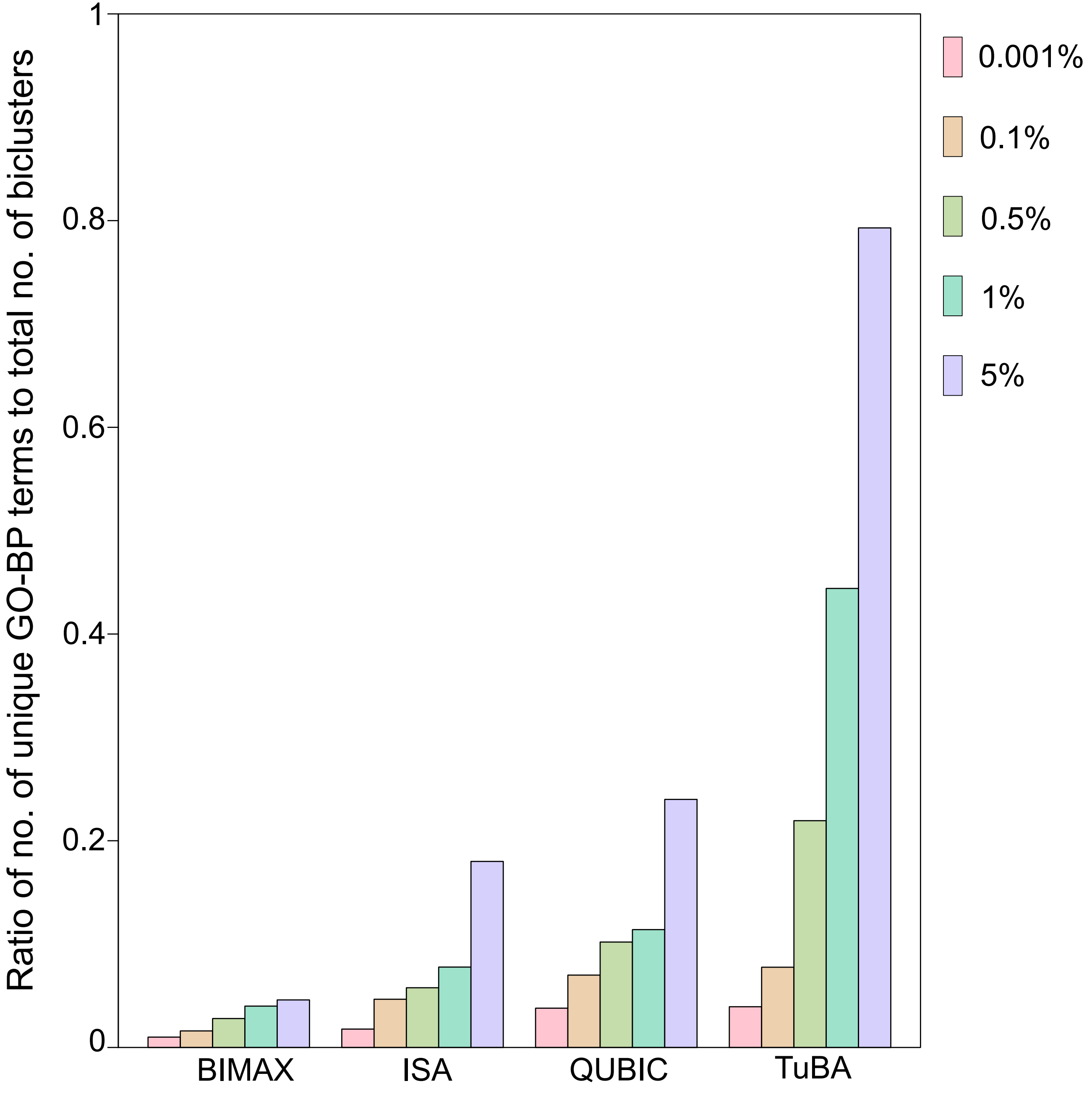

Ratio of no. of unique GO-BP terms to total no. of biclusters
1
0.8
0.6
0.4
0.2
0
BIMAX
ISA
QUBIC
TuBA
0.001%
0.1%
0.5%
1%
5%

Fig. S9A

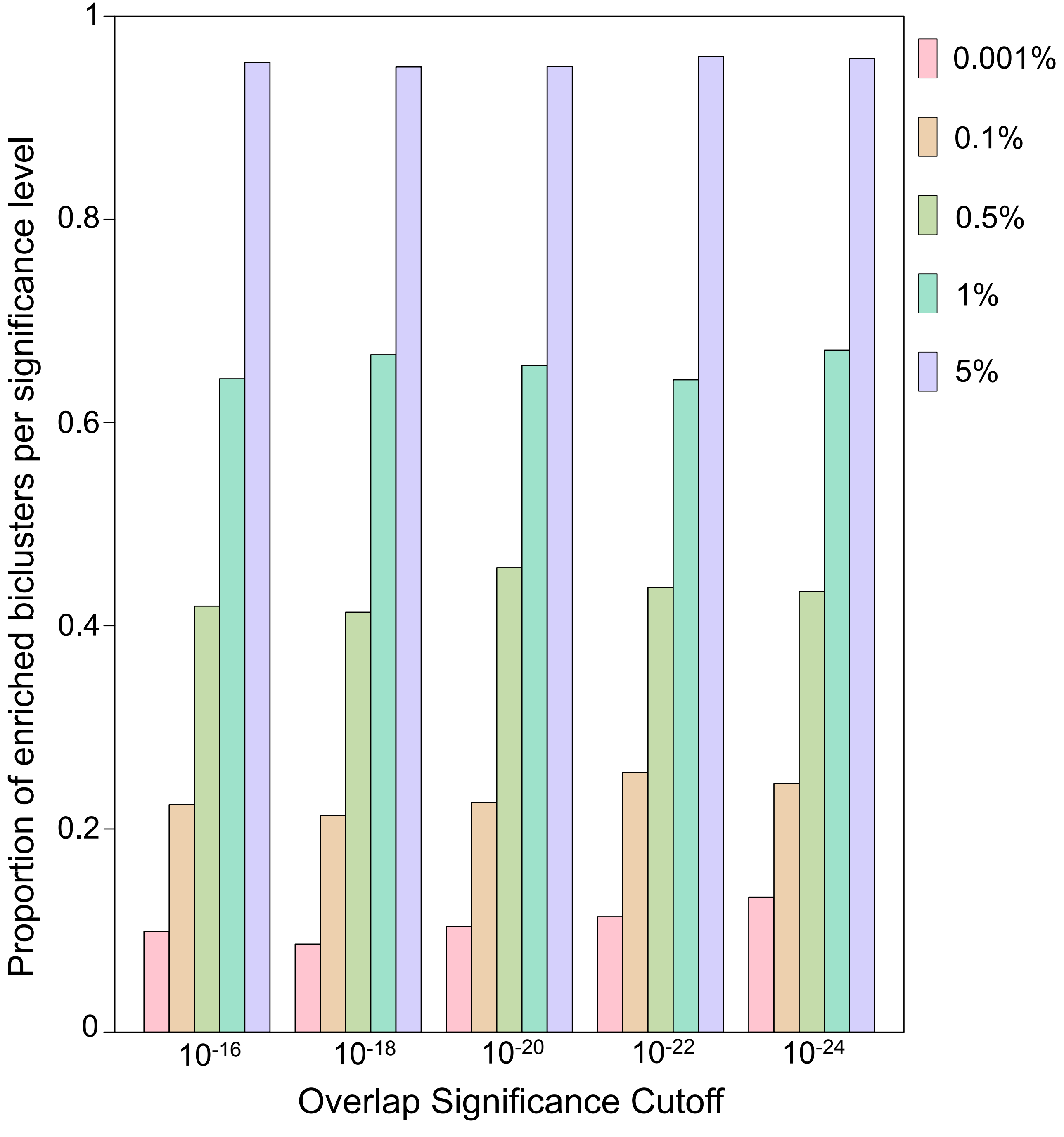

0.001%
0.1%
0.5%
1%
5%
Proportion of enriched biclusters per significance level
1
0.8
0.6
0.4
0.2
0
$10^{-16}$
$10^{-18}$
$10^{-20}$
$10^{-22}$
$10^{-24}$
Overlap Significance Cutoff

Fig. S9B

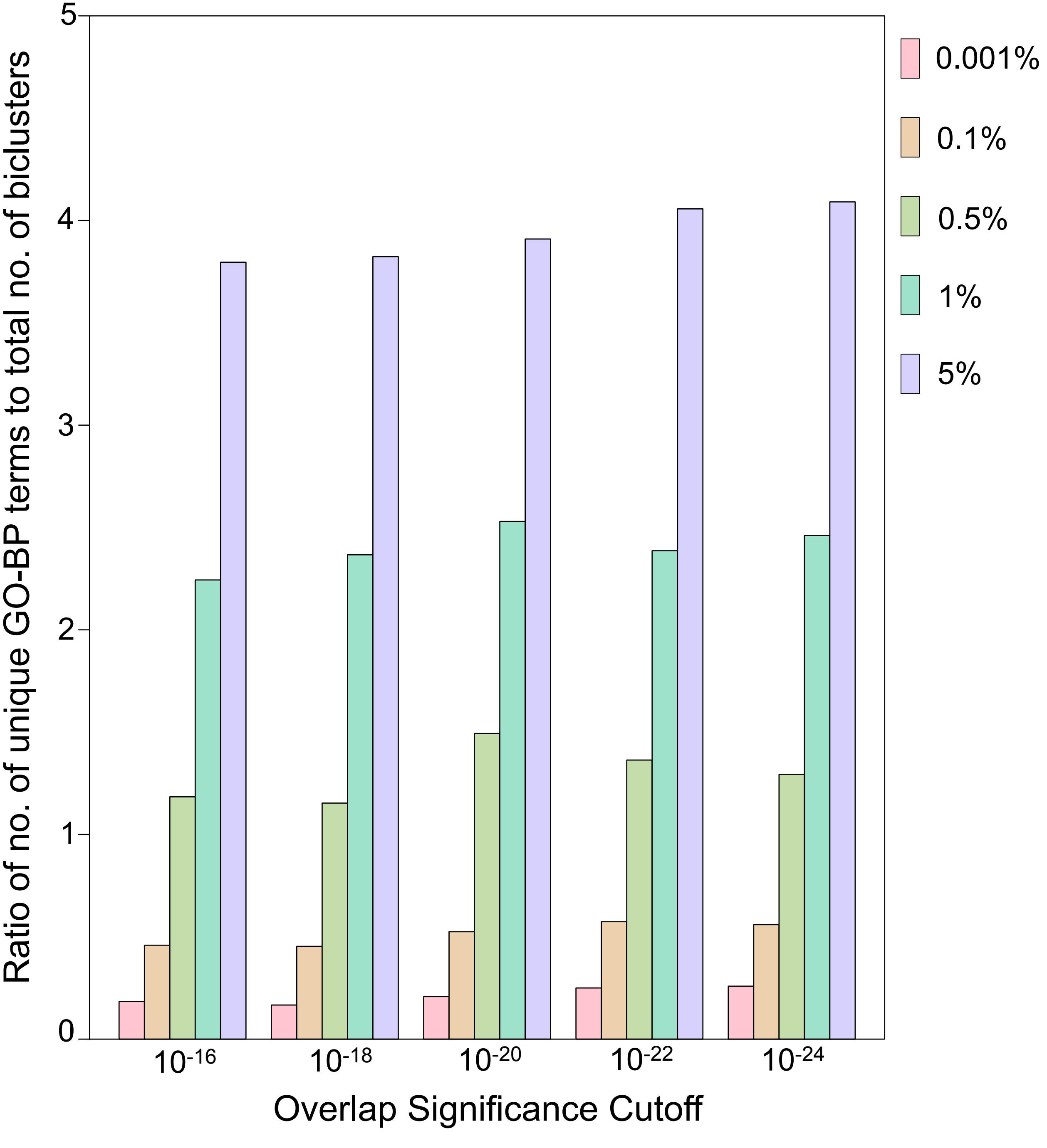

5
4
3
2
1
0
Ratio of no. of unique GO-BP terms to total no. of biclusters
0.001%
0.1%
0.5%
1%
5%
10-16
10-18
10-20
10-22
10-24
Overlap Significance Cutoff
