## Supplementary Methods for "TuBA: Tunable Biclustering Algorithm Reveals Clinically Relevant Tumor Transcriptional Profiles in Breast Cancer"

**Computation of significance values of overlaps for high expression**

We estimate the significance values for the pairwise overlaps using Fisher’s exact test. The setup of the contingency table for the test is illustrated with the help of **Fig. 1**. The rectangular box (containing both the circles) represents the set of all samples that have non-zero expression values for both gene 1 and gene 2. The red circle represents the set of top (5%, 10%, etc.) percentile samples for gene 1 or the set of all non-zero samples for gene 1, whichever is smaller, the blue circle represents the set of upper percentile samples for gene 2 or the set of all non-zero samples for gene 2, whichever is smaller.

In the Venn diagram (**Fig. 1B**), the regions labeled by a, b, c and d respectively represent:

**a:** Set of samples that are found in the upper percentile (or non-zero) sets of both gene 1 and gene 2

**b:** Set of samples that are found only in the upper percentile (or non-zero) set of gene 1

**c:** Set of samples that are found only in the upper percentile (or non-zero) set of gene 2

**d:** Set of samples that are neither found in the upper percentile (or non-zero) set of gene 1 nor in the percentile (or non-zero) set of gene 2

We use Fisher’s Exact test to calculate the probability of the null hypothesis that the odds ratio equals one (or equivalently, the independence of the rows and columns of the contingency table), against the alternative hypothesis that the odds ratio is > 1.

Since we do multiple hypotheses testing to compute the p-values for each pair of genes in our dataset, we need to correct the *p*-values for false discovery. We adjust the *p*-values using the Benjamini-Hochberg method.

**Computation of significance values of overlaps for low expression (RNASeq)**

For the case of low expression, the rectangular box (containing both the circles) represents the set of all samples in the dataset. The red circle represents the set of bottom (5%,10%, etc.) percentile samples for gene 1 *or* the set of all samples with zero expression for gene 1, whichever is larger, the blue circle represents the set of bottom (5%, 10% etc) percentile samples for gene 2 *or* the set of all samples with zero expression for gene 2, whichever is larger.

In the Venn diagram, the regions labeled by a, b, c and d respectively represent:

**a:** Set of samples that are found in the lower percentile (or zero expression) sets of both gene 1 and gene 2

**b:** Set of samples that are found only in the lower percentile (or zero expression) set of gene 1

**c:** Set of samples that are found only in the lower percentile (or zero expression) set of gene 2

**d:** Set of samples that are neither found in the lower percentile (or zero expression) set of gene 1 nor in the percentile (or zero expression) set of gene 2

**Permutation Test**

How likely is it for gene-pair associations to be identified in our graphs purely due to chance? We decided to investigate this question by performing a permutation test on the METABRIC dataset (1970 samples) with upper percentile set size cutoff: 5%. For each gene, we permuted the labels of the samples prior to ascertainment of the samples that corresponded to the top 5% respectively. The significance values for overlaps between every pair of genes were computed using the Fisher’s exact test. The histogram for the distribution of the p-values is shown in **Fig. S1A**. After adjusting for multiple hypotheses testing, none of the gene-pair p-values were found to be significant (**Fig. S1B**). We performed 100 such iterations, and did not identify a single significant gene-pair association in any of them.

**Robustness of TuBA’s biclusters**

We investigated the robustness of TuBA results against varying choices of its parameters by inquiring whether there was agreement between biclusters obtained for a given dataset. As our reference, we chose the biclusters obtained for TCGA (high expression) with the following choices for the parameters: (i) Percentile Cutoff: 5%, and (ii) Overlap Significance Cutoff: 1e-16 (The results for TCGA discussed in this paper correspond to the biclusters obtained for these set of parameter values). Subsequently, we applied TuBA to the TCGA dataset with larger cutoffs for the size of the top percentile sets - 7.5%, 10%, 12.5%, 15%. Corresponding to each of these choices of percentile cutoff, we also made different choices for the overlap significance cutoff based on our proposed heuristic (**Supplementary Table 4**). Pairwise comparisons between our reference biclusters, and the biclusters obtained with the other sets of parameter choices, identified those reference biclusters that shared a significant proportion of their genes and samples with biclusters corresponding to the other sets of parameter choices. We observed that 95% of our reference biclusters (334 out of 353 biclusters) were enriched (FDR < 0.001) in at least one bicluster in the set of biclusters corresponding to the percentile cutoff of 7.5%, and overlap significance cutoff of 1e-19. The agreement decreases as we increase the size of the percentile sets - we observed that 86% of the reference biclusters (304 out of 353 biclusters) were enriched (FDR < 0.001) in at least one bicluster in the set of biclusters corresponding to the percentile cutoff of 15%, and overlap significance cutoff of 1e-30. There was a significant difference (Mann-Whitney U-test *p* < 1e-05) in the number of genes contained in biclusters that matched, compared to the ones that did not match; while the median size of biclusters that matched was 20 (range: 3–1012), the median size of biclusters that did not match was 4 (range: 3–60). Thus, we can conclude that some of the smaller biclusters that correspond to alterations/dyregulation in small subsets of tumors were not identified as we increased the size of the percentile sets. However, we were able to validate most of our reference biclusters in biclusters obtained with other choices of the parameters, thus demonstrating their robustness.

Another aspect that we investigated was the level of agreement between the reference biclusters and the biclusters obtained by varying the overlap significance cutoff. Our reference biclusters once again corresponded to the ones obtained for TCGA (high expression) with the following choices for the parameters: (i) Percentile Cutoff: 5%, and (ii) Overlap Significance Cutoff: 1e-16. We compared these biclusters to the ones obtained corresponding to the percentile cutoff of 10% and overlap significance cutoff values ranging between 1e-23 to 1e-28 (in increments of negative powers of 10) (**Supplementary Table 3**). For the overlap significance cutoff of 1e-23, we observed that 93% (327 out of 353 biclusters) of the reference biclusters were enriched (FDR < 0.001) in at least one bicluster in the set of biclusters corresponding to the percentile cutoff of 10%. On the other hand, the agreement decreased to 81% (288 out of 353 biclusters) as we increased the level of overlap significance to 1e-28. This is reasonable since we decrease the number of gene-pairs in our graph as we increase the level of overlap significance. This leads to fewer genes and samples in the graphs overall, and an omission of gene-pair associations that correspond to some of our reference biclusters. However, despite a five-fold difference in the significance level of overlap there is still good agreement between the reference biclusters and the biclusters obtained with other choices of the overlap significance cutoff.

**Association with copy number gain (METABRIC & TCGA):**

For the biclusters obtained in the high expression case it is useful to identify the underlying reason for the relatively high expression of the genes in the bicluster. The dominant alteration in many tumor types is gain in copy number that results in higher levels of the corresponding transcripts. We adopt the following procedure in order to identify whether gain in copy number might be the underlying mechanism for some of our biclusters:

1. List the genes that constitute the bicluster
2. Pick a gene from the list prepared above
3. List all the samples that have a copy number gain (> 0) for the chosen gene and are also present in the gene expression dataset
4. List all the samples that are in the top percentile set for the given gene and have copy number data available

We employ Fisher’s Exact test to look for enrichment of samples with copy number gains for the given gene in the bicluster. We refer to the Venn diagram above, where now for the copy number analysis the rectangular box (containing both the circles) represents the set of all the samples that have both copy number and gene expression data available, the red circle represents the set of top percentile samples corresponding to the given gene present in the bicluster, and the blue circle represents the set of samples that have a gain in copy number for the given gene.

In the Venn diagram, the regions labeled by a, b, c and d respectively represent:

**a:** Set of top percentile samples in the bicluster that also have a gain in copy number for the given gene

**b:** Set of top percentile samples in the bicluster that do not have a copy number gain for the given gene

**c:** Set of samples that have a gain in copy number for the given gene but are not present in the bicluster

**d:** Set of samples that are neither present in the bicluster nor do they have a gain in copy number for the given gene

We prepare such 2×2 contingency tables for each gene in every bicluster to compute the *p*-value of the null hypothesis that the odds ratio equals one, against the alternative hypothesis that the odds ratio is greater than 1. We adjust the *p*-values using the Benjamini-Hochberg method for false discovery.

**Association with copy number deletion (TCGA - RNASeq):**

We adopt the following procedure to identify whether loss in copy number (deletions) are associated with some biclusters:

1. List the genes that constitute the bicluster
2. Pick a gene from the list prepared above
3. List all the samples that have a copy number deletion (< 0) for the chosen gene and are also present in the gene expression dataset
4. List all the samples that are in the bottom percentile set for the given gene and have copy number data available

In the Venn diagram then, the rectangular box (containing both the circles) represents the set of all samples that have both copy number and gene expression data available. The red circle represents the set of bottom percentile samples corresponding to the given gene in the bicluster, the red circle represents the set of samples that have copy number deletion for the given gene.

In the Venn diagram, the regions labeled by a, b, c and d respectively represent:

**a:** Set of bottom percentile samples in the bicluster that also have deletion in copy number of the given gene

**b:** Set of bottom percentile samples in the bicluster that do not have copy number deletion for the given gene

**c:** Set of samples that have copy number deletion for the given gene but are not present in the bicluster

**d:** Set of samples that are neither present in the bicluster nor do they have copy number deletion for the given gene

As for the case for copy number gain, 2×2 contingency tables are prepared for each gene in every bicluster to compute the *p*-values that are then adjusted by the Benjamini-Hochberg method for false discovery. These individual *p*-values are then combined for each bicluster to arrive at a singular p-value by using the Fisher method.

**Subtype enrichment of biclusters (ER/HER2)**

For METABRIC, we used the ER & HER2 status available in the clinical file based on their expression levels to identify the ER/HER2 subtype for the sample. For GEO, we used the expression levels of the transcripts to identify the subtype for each sample. For TCGA as well, we used the expression levels of the transcripts of ER and HER2 to classify the samples into the four subtypes.

The setup of the test to find enrichment for the ER/HER2 subtypes is similar to the one described above for copy number gain. We can take the help of the Venn diagram once again to illustrate how the contingency tables for Fisher’s Exact test are prepared. In the Venn diagram, the rectangular box (containing both the circles) represents the set of all samples that have been assigned an ER/HER2 subtype. The red circle represents the set of all the samples corresponding to a bicluster, the blue circle represents the set of samples that belong to a given subtype.

In the Venn diagram, the regions labeled by a, b, c and d respectively represent:

**a:** Set of samples in the bicluster that also belong to the given ER/HER2 subtype

**b:** Set of samples in the bicluster that do not belong to the given ER/HER2 subtype

**c:** Set of samples for the given subtype that are not present in the bicluster

**d:** Set of samples that are neither present in the bicluster, nor are they of the given subtype

For each of the four ER/HER2 subtypes, we prepare such contingency tables for every bicluster and check whether some of the biclusters are enriched in specific subtypes.

**Subtype enrichment of biclusters (PAM50)**

For METABRIC, we used the PAM50 subtype calls available in the clinical file, while for TCGA we used the PAM50 calls available for a subset of samples in the clinical file from the 2012 Nature study. For both the datasets, we identified the samples that had been assigned to at least one of the five subtypes (Basal, Her2, Luminal A, Luminal B, Normal-like) and used our algorithm to identify the biclusters for the dataset comprising only these samples.

Similar to the case above, in the Venn diagram the rectangular box (containing both the circles) represents the set of all samples that have an unambiguous PAM50 subtype assignment available. The red circle represents the set of all the samples corresponding to a bicluster, the blue circle represents the set of samples that belong to a given subtype.

In the Venn diagram, the regions labeled by a, b, c and d respectively represent:

**a:** Set of samples in the bicluster that also belong to the given PAM50 subtype

**b:** Set of samples in the bicluster that do not belong to the given PAM50 subtype

**c:** Set of samples for the given subtype that are not present in the bicluster

**d:** Set of samples that are neither present in the bicluster nor are they of the given subtype

For each of the five PAM50 subtypes, we prepare such contingency tables for every bicluster and check for enrichment of biclusters in every subtype.

**Immune cell infiltration and stromal cell signatures**

We stratified the samples for the TCGA RFS dataset based on the ESTIMATE scores for the level of stromal cells present, and the infiltration level of immune cells in tumor tissues into three groups: (i) top 25 percentile, (ii) intermediate 50 percentile, and (iii) bottom 25 percentile.

In the Venn diagram, the rectangular box (containing both the circles) represents the set of all samples in the gene expression dataset that have ESTIMATE scores for stromal cells and immune cell infiltration. The red circle represents the set of all the samples corresponding to a bicluster, the blue circle represents the set of samples that belong to group (i) for each case (stromal and immune respectively).

In the Venn diagram, the regions labeled by a, b, c and d respectively represent:

**a:** Set of samples in the bicluster that also belong to group (i)

**b:** Set of samples in the bicluster that do not belong to group (i)

**c:** Set of samples in group (i) that are not present in the bicluster

**GTEx Gene Pair Associations Enrichment:**

For the GTEx breast tissue dataset, we compared the upper 10 percentile samples between all those pairs of genes that are also present in the TCGA dataset and calculated the significance of overlaps to obtain a matrix of *p-*values for all pairs of genes. For each bicluster, we listed all the gene pairs and identified the number of gene pairs that were found to have an overlap significance above a predetermined value (*p* ≤ 10^-05^)^.^ We then set up contingency tables for Fisher’s Exact test for each bicluster to test for the significance of the proportion of gene pairs that are enriched in GTEx.

In the Venn diagram, the rectangular box (containing both the circles) represents the total set of all the gene pairs present in biclusters from TCGA. The red circle represents the set of all gene pairs corresponding to a bicluster from TCGA, the blue circle represents the set of gene pairs that are enriched in GTEx.

In the Venn diagram, the regions labeled by a, b, c and d respectively represent:

**a:** Set of gene pairs present in bicluster as well as enriched in GTEx

**b:** Set of gene pairs present present in bicluster from TCGA but not enriched in GTEx

**c:** Set of gene pairs enriched in GTEx but not present in bicluster

**d:** Set of gene pairs not enriched in GTEx and not present in bicluster

**Impact of the choice of TuBA’s knobs**

We illustrate how the choices of the parameters determine the final biclusters with the help of two examples from the application of TuBA to the 908 samples from the TCGA RFS dataset. If we choose a 5% upper percentile cutoff and an overlap significance cutoff of *p* ≤ 10^-16^, one of the biclusters we obtain comprises solely of genes from the Cancer-Testis Antigen family: MAGEA2, MAGEA3, MAGEA6, MAGEA10, CSAG1, CSAG2. However, this bicluster is not identified for percentile set cutoff choices of upper 10% and 20% with the overlap significance cutoff choices of *p* ≤ 10^-25^ and *p* ≤ 10^-36^ respectively. This suggests that these genes are only expressed in ≤ 5% of all BRCA tumors. Thus, increase in percentile cutoff results in a systematic omission of aberrant signatures found in comparatively small subsets of the population, such as this one. In contrast, we find another small bicluster comprising genes from the Cancer-Testis Antigen family - CTAGE4, CTAGE6, CTAGE9 - for all three upper percentile cutoffs (5%, 10% and 20%) and the respective overlap cutoffs, suggesting that these genes are aberrantly expressed in a larger proportion of the population (at least 10%).

The second example is that of the bicluster corresponding to the Her2 amplicon. **Fig. S2A** shows the number of patients/samples present in the bicluster for the upper percentile cutoff of 5%, as the overlap cutoff is lowered from 10^-20^ to 10^-16^. We notice that even though there is a reduction in the significance level of overlap by four orders of magnitude, no new samples get added. We can infer that the choice of the upper percentile cutoff at 5% put a cap on the maximum number of samples that may be present in the bicluster; lowering of the overlap cutoff does not lead to an increase in the number of samples in the bicluster because almost all of them were already identified at the higher significance level of 10^-20^. When the upper percentile cutoff is chosen at 10%, we initially observe an increase in the number of samples in the bicluster as we lower the overlap cutoff, however the number of patients added to the biclusters with further lowering of the overlap cutoff begins to decrease as we approach *p* = 10^-23^ (**Fig. S2B**). Further reduction in the overlap cutoff results in the addition of only a handful of samples to the bicluster, but is accompanied by a substantial increase in the number of edges in the overall graph.

**Degrees of genes as indicators of their importance**

For biclusters associated with gain in copy number, we can infer the extent as well as the focal nature of copy number gains based on the degrees of the constituent genes. We illustrate this with the example of the HER2 amplicon, which is a robust bicluster in all three datasets. **Fig. S6A**, **S6B**, and **S6C** show the degrees of the genes that comprise the HER2 bicluster in TCGA, METABRIC, and GEO respectively. The genes are arranged in increasing order of their start site positions on the chromosome (Chr17). Higher degrees of specific genes suggest focal gains in copy number at multiple sites along the chromosomal segment – one around CDK12-STARD3, one around ERBB2, and another one around MSL1-CASC3. This is a simplistic guess, since there are additional layers of transcriptional regulation that may modulate the expression levels of these genes. Nonetheless, given the profile of the degrees of the genes, it would be prudent to investigate the role of genes with higher degrees in tumor progression/development. For our example, CDK12 and ERBB2 would definitely qualify as candidates for further investigation.

**Comparison with other biclustering methods**

We applied TuBA to the DLBCL dataset, and performed a GO-BP enrichment analysis on the *seeds* of the resulting biclusters using GeneSCF. TuBA discovered 94 biclusters in total (45 biclusters for high, and 49 biclusters for low expression) (**Supplementary Table 7**), DeBi discovered 127 biclusters in total (68 biclusters with up-regulated genes, and 59 biclusters with down-regulated genes), ISA discovered 49 biclusters, OPSM discovered just 12 biclusters, QUBIC discovered 100 biclusters (the default number), and SAMBA discovered 128 biclusters. **Fig. 8A** shows the proportions of GO-BP enriched biclusters for five different significance levels (FDRs) – 0.001%, 0.1%, 0.5%, 1%, and 5%. For the FDR cutoff of less than 5%, almost all the biclusters for every biclustering algorithm were enriched in at least one GO-BP term. TuBA had 2 non-enriched biclusters out of 94 biclusters, SAMBA had 2 non-enriched biclusters out of 128 biclusters, and QUBIC had 1 bicluster out of 100 that was not enriched in a GO-BP term. We decided to investigate how many of the biclusters identified by other algorithms did not share genes between them. For this, we filtered the biclusters to exclude the ones that have significant overlaps between their genes (i.e. biclusters with hypergeometric test FDR < 0.001). DeBi had only 9 biclusters, out of a total 127 biclusters, that did not have significant overlaps of their genes with any other bicluster (7 biclusters with up-regulated genes, and 2 biclusters with down-regulated genes), ISA had 15 biclusters, out of a total 49 biclusters, that did not share genes, OPSM had 3 biclusters, out of a total of 12 biclusters, that did not share genes, QUBIC had 29 biclusters out of 100 biclusters, that did not share genes, and SAMBA had just 10 biclusters, out of a total of 128 biclusters, that did not share genes. Since most of the biclusters discovered by these algorithms shared genes with other biclusters, we expected redundancy in the enriched GO-BP terms. We identified the top 5 GO-BP terms for every bicluster obtained by each algorithm (not every bicluster was enriched in 5 distinct GO-BP terms, some had less than 5, while others were not enriched in any term). We then prepared a list comprising all the unique GO-BP terms for the complete set of biclusters for a given algorithm. For the five levels of significance of enrichment, TuBA identified sets comprising 337, 218, 98, 51, and 25 distinct GO-BP terms, respectively. On the other hand, DeBi identified sets comprising 259, 146, 120, 68, and 34 distinct GO-BP terms, ISA identified sets with 172, 100, 69, 39, and 25 distinct GO-BP terms, OPSM identified sets with 41, 26, 24, 13, and 8 distinct GO-BP terms, QUBIC identified sets with 237, 172, 106, 54, and 22 distinct GO-BP terms, while SAMBA identified sets with 250, 138, 120, 72, and 37 distinct GO-BP terms, respectively. **Fig. 8C** shows the ratios of the number of elements in these sets to the total number of biclusters for each algorithm, for the five different significance levels.

Apart from the DLBCL dataset, we also investigated the TCGA dataset with the following biclustering algorithms: (i) BIMAX, (ii) ISA, (iii) QUBIC, and (iv) SAMBA. We used the respective default parameters for all four biclustering algorithms. TuBA discovered 556 biclusters in total (353 biclusters for high, and 203 biclusters for low expression), BIMAX discovered 100 biclusters (default), ISA discovered 244 biclusters, QUBIC discovered 100 biclusters (default), and SAMBA discovered 405 biclusters (**Supplementary Table 5**). **Fig. 8B** shows the proportion of GO-BP terms enriched biclusters of each algorithm for five different significance levels. This time, when we filtered the biclusters obtained from the other algorithms to exclude the ones that have significant overlaps between genes (i.e. biclusters with hypergeometric test FDR < 0.001), we discovered that none of the four algorithms identified a single bicluster that did not have significant overlap of its genes with at least one other bicluster.

Once again, we identified unique sets of GO-BP terms for the results of each biclustering algorithm. For the five levels of significance of enrichment, TuBA identified unique sets with 1874, 1099, 556, 220, and 99 distinct GO-BP terms, respectively. In sharp contrast, BIMAX identified sets with just 23, 17, 12, 7, and 5 GO-BP terms, ISA identified sets with 148, 148, 51, 36, and 24 distinct GO-BP terms, QUBIC identified sets with 174, 56, 32, 24, and 4 distinct GO-BP terms, while SAMBA identified sets with 490, 155, 72, 34, and 11 distinct GO-BP terms, respectively. **Fig. 8D** shows the ratios of the number of elements in these sets to the total number of biclusters for each algorithm, for the five different significance levels.

Finally, we investigated the investigated the METABRIC dataset with the following biclustering algorithms (apart from TuBA): (i) BIMAX (ii) ISA, and (iii) QUBIC (We could not get an output for SAMBA from Expander as the METABRIC dataset exceeded the memory limit defined by the software). We used the respective default parameters for all three biclustering algorithms. TuBA discovered 340 biclusters (high expression), BIMAX discovered 100 biclusters (default), ISA discovered 90 biclusters, and QUBIC discovered 100 biclusters (default) (**Supplementary Table 5**). **Fig. S8A** shows the proportion of GO-BP terms enriched biclusters of each algorithm for five different significance levels. When we filtered the biclusters obtained from the other algorithms to exclude the ones that have significant overlaps between genes (i.e. biclusters with hypergeometric test FDR < 0.001), we discovered that none of the three algorithms identified a single bicluster that did not have significant overlap of its genes with at least one other bicluster. We identified unique sets of GO-BP terms for the results of each biclustering algorithm. For the five levels of significance of enrichment, TuBA identified unique sets with 1348, 755, 373, 132, and 67 distinct GO-BP terms, respectively. BIMAX identified sets with just 23, 20, 14, 8, and 5 GO-BP terms, ISA identified sets with 81, 35, 26, 21, and 8 distinct GO-BP terms, while QUBIC identified sets with 120, 57, 51, 35, and 19 distinct GO-BP terms, respectively. **Fig. S8B** shows the ratios of the number of elements in these sets to the total number of biclusters for each algorithm, for the five different significance levels.

**GO Term Enrichment Across Different Choices of Overlap Cutoff**

We looked at GO-BP-term enrichments for TuBA’s biclusters for TCGA for five different choices of the overlap significance cutoffs - 10^-16^, 10^-18^, 10^-20^, 10^-22^, 10^-24^. For these five choices of the overlap cutoff the number of biclusters discovered by TuBA were – 353, 300, 221, 176, and 143, respectively. Although, the total number of biclusters obtained differed for each choice, the proportion of enriched biclusters at different significance levels remained similar irrespective of the parameter choice (**Fig. S9A**). Similarly, the ratios of the number of unique GO-BP terms to the total number of biclusters were consistent across all five choices of the overlap cutoffs (**Fig. S9B)**.

**Pseudocode for TuBA**

READ gene expression file

READ user input for size of percentile sets

READ user specification for whether the percentile sets should correspond to samples with highest or lowest gene expression values

FOR each gene

SORT samples in increasing order of their expression values

LIST samples that lie in the upper-most or lower-most percentile set

ENDFOR

FOR every pair of genes

COMPUTE number of shared samples between the respective percentile sets

PREPARE 2×2 contingency table such that the first row corresponds to samples in percentile set 1, the second row corresponds to samples in the dataset not present in percentile set 1, the first column corresponds to samples present in percentile set 2, while the second column corresponds to samples in the dataset not present in percentile set 2.

COMPUTE the probability of the null hypothesis (p-value) that the odds ratio equals one (or equivalently, the independence of the rows and columns of the contingency table), against the alternative hypothesis that the odds ratio is greater than 1.

ENDFOR

CORRECT the p-values for multiple hypotheses testing using the Benjamini-Hochberg method

LIST all gene-pairs that have a corrected p-value less than 0.05

READ user input for p-value cutoff for overlapping samples

LIST all gene-pairs that have p-values less than or equal to the user-specified cutoff

GENERATE undirected graph showing all these gene-pairs as nodes linked together by edges

REMOVE all nodes with degree 1 (nodes with one edge)

REMOVE all nodes linked with two nodes that are not connected with each other (to ensure that the elementary units in our graphs are triangles)

COMPUTE degrees of all nodes (genes) in graphs

SET Node_With_Highest_Degree = Node with the highest degree

Cluster_Serial_Number = 1

WHILE Node_With_Highest_Degree is non-null

Associated_Nodes = All nodes that are linked to Node_With_Highest_Degree

WHILE length of Associated_Nodes is non-zero

FOR all the nodes in Associated_Nodes

LIST all nodes that are linked to Associated_Nodes

ENDFOR

ASSIGN Cluster_Serial_Number to all nodes in the union of all these sets of nodes, as well as to the Node_With_Highest_Degree

ENDWHILE

INCREMENT Cluster_Serial_Number by 1

REMOVE all the nodes and links that contain the nodes identified in cluster

REMOVE all genes identified in cluster, from the list that contains their degrees in the graph

SET Node_With_Highest_Degree = Node with highest degree in the remaining list

ENDWHILE

SET Cluster_Number = 1

SET Nodes_In_Cluster = Genes in cluster number 1

SET Index_Number = 0

SET Size_of_Largest_Clique = 1

WHILE Index_Number is less than the length of Nodes_In_Cluster

IF Size_of_Largest_Clique is less than 3 OR there are no gene-pairs left THEN

INCREMENT Index_Number by 1

EXTRACT the cluster (subgraph) that contains all the links in graph that exclusively connect only those nodes whose Cluster_Serial_Number matches the Index_Number

ENDIF

COMPUTE the largest cliques present in cluster using the Bron-Kerbosch algorithm

IF multiple cliques qualify as largest THEN

SET Nodes_of_Seed = Union of the set of nodes of the subset of these cliques that have non-zero intersection

ELSE

SET Nodes_of_Seed = Nodes present in largest clique

ENDIF

REMOVE all links in the cluster that connect to nodes in Nodes_of_Seed

COMPUTE largest clique in reduced graph

SET Size_of_Largest_Clique = Number of nodes in largest clique

ENDWHILE

GENERATE undirected graph showing all the gene-pairs that have p-values less than or equal to the user specified cutoff

FOR all sets of Nodes_of_Seed

LIST Nodes_To_Be_Added = Nodes that have links with at least two nodes in Nodes_of_Seeds

SET Genes_of_Bicluster = Union of Nodes_To_Be_Added and Nodes_of_Seeds

ENDFOR

FOR all sets of Genes_of_Bicluster

LIST gene-pairs present in bicluster

SET Samples_of_Bicluster  = Union of the sets of shared samples in percentile sets of listed gene-pairs

SET Degrees_of_Gene = Number of links that connect genes in bicluster to other genes within the bicluster

ENDFOR

**SUPPLEMENTARY FIGURE LEGENDS**

**Fig. S1.** (A) Histogram for overlap significance values (in –log_10_ scale) based on a permutation test on the METABRIC dataset. The p-values were not corrected for multiple hypotheses testing. (B) Histogram for overlap significance values (in –log_10_ scale) after correcting for multiple hypotheses testing.

**Fig. S2.** Impact of the size of top percentile set for the bicluster corresponding to the HER2 amplicon (17q12). (A) Number of samples in bicluster as the overlap significance is lowered from 10^-20^ to 10^-16^ for percentile set size of 5%, (B) number of samples in bicluster as the overlap significance is lowered from 10^-35^ to 10^-23^ for percentile set size of 10%.

**Fig. S3.** Enrichment of biclusters consisting of proximally located genes with copy number gains in the PAM50 subtypes for (A) METABRIC and (C)(E) TCGA, respectively. The biclusters are represented by horizontal bars in each panel, color-coded according to the chromosome number of their constituent genes. Panels (B),(D) and (F) show the remaining biclusters arranged according to their serial numbers in **Supplementary Table 3** for METABRIC and TCGA, respectively. The ones that are associated with copy number (CN) gains of genes located at distant chromosomal sites are shown in red while those associated with loss are shown in green. The rest are shown in black. Note, the thickness of the bar in each figure depends on the total number of biclusters displayed in that figure and so does not represent its chromosomal extent.

**Fig. S4.** (A) Enrichment of biclusters from TCGA consisting of proximally located genes with copy number loss in the ER/HER2 subtypes. The biclusters are represented by horizontal bars in each panel, color-coded according to the chromosome number of their constituent genes. Panel (B) shows the remaining biclusters arranged according to their serial numbers in **Supplementary Table 3**. The ones that exhibit copy number loss of genes located at distant chromosomal sites are shown in green, while the rest are shown in black. Note, the thickness of the bar in each figure depends on the total number of biclusters displayed in that figure and so does not represent its chromosomal extent.

**Fig. S5.** Enrichment of biclusters from GEO in the subtypes based on ER and HER2 status. The biclusters have been arranged according to their serial numbers in **Supplementary Table 3** (Bicluster 1 on top) and are represented by horizontal black bars.

**Fig. S6.** Degrees of genes in the bicluster corresponding to the HER2 amplicon (17q12) for (A) TCGA, (B) METABRIC and (C) GEO respectively.

**Fig. S7.** (A) Kaplan-Meier survival curve for the set of patients in the Her2 (17q12) bicluster (red) compared to the remaining set of patients (blue) for the METABRIC dataset, and (B) the graph corresponding to the bicluster.

**Fig. S8.** (A) Proportions of GO-BP terms enriched biclusters for each biclustering method at five different significance levels for the METABRIC dataset. (B) Ratios of number of unique GO-BP terms and total number of biclusters at five different significance levels for the METABRIC dataset.

**Fig. S9.** (A) Proportions of GO-BP terms enriched biclusters obtained by TuBA at five different significance levels for five choices of the overlap significance cutoff for the TCGA dataset. (B) Ratios of number of unique GO-BP terms to the total number of biclusters at five different significance levels for five choices of the overlap significance cutoff for the TCGA dataset.
